# Identification of SUMO targets required to maintain human stem cells in the pluripotent state

**DOI:** 10.1101/2020.12.22.423944

**Authors:** Barbara Mojsa, Michael H. Tatham, Lindsay Davidson, Magda Liczmanska, Emma Branigan, Ronald T. Hay

## Abstract

To determine the role of SUMO modification in the maintenance of pluripotent stem cells we used ML792, a potent and selective inhibitor of SUMO Activating Enzyme. Treatment of human induced pluripotent stem cells with ML792 initiated changes associated with loss of pluripotency such as reduced expression of key pluripotency markers. To identify putative effector proteins and establish sites of SUMO modification, cells were engineered to stably express either SUMO1 or SUMO2 with TGG to KGG mutations that facilitate GlyGly-K peptide immunoprecipitation and identification. A total of 976 SUMO sites were identified in 427 proteins. STRING enrichment created 3 networks of proteins with functions in regulation of gene expression, ribosome biogenesis and RNA splicing, although the latter two categories represented only 5% of the total GGK peptide intensity. The remainder have roles in transcription and chromatin structure regulation. Many of the most heavily SUMOylated proteins form a network of zinc-finger transcription factors centred on TRIM28 and associated with silencing of retroviral elements. At the level of whole proteins there was only limited evidence for SUMO paralogue-specific modification, although at the site level there appears to be a preference for SUMO2 modification over SUMO1 in acidic domains. We show that SUMO is involved in the maintenance of the pluripotent state in hiPSCs and identify many chromatin-associated proteins as bona fide SUMO substrates in human induced pluripotent stem cells.

## Introduction

Pluripotent cells display the property of self-renewal and have the capacity to generate all of the different cells required for the development of the adult organism. The pluripotent state is defined by a specific gene expression programme driven by expression of the core transcription factors OCT4, SOX2 and NANOG that sustain their own expression by virtue of a positive, linked autoregulatory loop, while activating genes required to maintain the pluripotent state and repressing expression of the transcription factors for lineage specific differentiation (1). Once terminally differentiated, somatic cell states are remarkably stable, however, forced expression of key pluripotency transcription factors that are highly expressed in embryonic stem cells (ESCs), including OCT4, SOX2 and NANOG, leads to reprogramming back to the pluripotent state (2–4). Under normal circumstances the efficiency of reprogramming is very low and it is clear that there are roadblocks to reprogramming designed to safeguard cell fates (5, 6). The Small Ubiquitin like Modifier (SUMO) has emerged as one such roadblock and reduced SUMO expression decreases the time taken and increases the efficiency of reprogramming in mouse cells (7–9). Three SUMO paralogues, known as SUMO1, SUMO2 and SUMO3 are expressed in vertebrates. Based on almost indistinguishable functional and structural features SUMO2 and SUMO3 are collectively termed SUMO2/3 and share about 50% amino acid sequence identity with SUMO1. SUMO proteomics studies have revealed very high numbers of cellular SUMO substrates (see (10) for a review) and as a consequence SUMO can influence a wide range of biological processes. However, the proportion of the total the cellular pool of a protein that is SUMOylated varies greatly. There are rare examples of almost constitutively SUMOylated proteins such as the Ran GTPase RanGAP1 (11), although the majority of substrates have such low modification stoichiometry that identification from purified SUMO conjugates using Immuno-Blotting is close to the detection limit (for review see (12)). Conjugation of SUMO to protein substrates is mechanistically similar to that of ubiquitin conjugation but is carried out by a completely separate enzymatic pathway. SUMOs are initially translated as inactive precursors that require a precise proteolytic cleavage, carried out by a set of SUMO specific proteases (SENPs), to expose the terminal carboxyl group of a Gly-Gly sequence that ultimately forms an isopeptide bond with the ε−amino group of a lysine residue in the modified protein. The hetrodimeric E1 SUMO Activating Enzyme (SAE1/SAE2) uses ATP to adenylate the C-terminus of SUMO, before forming a thioester with a cysteine residue in a second active site of the enzyme and releasing AMP. SUMO then undergoes a transthioation reaction on to a cysteine residue in the only E2 SUMO conjugating enzyme Ubc9. Assisted by a small group of E3 SUMO ligases, including the PIAS proteins, RanBP2 and ZNF451 the SUMO is transferred directly from Ubc9 onto target proteins (13). Modification of target proteins may be short-lived, with SUMO being removed by a group of SENPs. Together this creates a highly dynamic SUMO cycle where the net SUMO modification status of proteins is determined by the rates of SUMO conjugation and deconjugation (12). Preferred sites of SUMO modification conform to the consensus ψKxE, where ψ represents a large hydrophobic residue (14, 15). A conjugation consensus is present in the N-terminal sequence of SUMO2 and SUMO3 and thus permits self-modification and the formation of SUMO2/3 chains (16). As a strict consensus is absent from SUMO1, it does not form chains as readily as SUMO2/3 (12). Once linked to target proteins SUMO allows the formation of new protein-protein interactions as the modification can be recognised by proteins containing a short stretch of hydrophobic amino acids termed a SUMO interaction motif (17).

Stem cell lines are an excellent model to study the mechanisms that control self-renewal and pluripotency. Mouse ESCs have been widely used for these studies as the cells can also be used in mice for in vivo studies. However, they display different characteristics from human ESCs. Mouse ESCs require leukemia inhibitory factor (LIF) and bone morphogenetic protein (BMP) signalling to maintain their self-renewal and pluripotency (18, 19). In contrast LIF does not support self-renewal and BMPs induce differentiation in human ESCs (20–22). The maintenance of the pluripotent state of hESCs requires basic fibroblast growth factor (bFGF, FGF2) and activin/nodal/TGF-b signalling along with inhibition of BMP signalling (23, 24). These differences may reflect the particular developmental stages at which ESC lines are established in vitro from mouse and human blastocysts, or may be due to differences in early embryonic development (25). As hESCs are derived from embryos their use is limited, but human Induced Pluripotent Stem Cells (hiPSC) can be derived by reprogramming normal somatic cells and display most of the characteristics of hESCs (2) and are now widely used to study self-renewal and pluripotency in humans.

To determine the role of SUMO modification in hiPSCs we made use of ML792, a highly potent and selective inhibitor of the SUMO Activating Enzyme (26). Treatment of hiPSCs with this inhibitor rapidly blocks SUMO modification allowing endogenous SUMO proteases to strip SUMO from targets. When used over the course of 48 hours hiPSCs treated with ML792 lose the majority of SUMO conjugation but show no large-scale changes to the cellular proteome nor loss of viability, although markers of pluripotency are reduced. Decreased expression of selected pluripotency markers seems to be a consequence of SUMO removal from key targets rather than large scale changes to the proteome. SUMO site proteomic analysis of hiPSCs reveals extensive SUMO modification of proteins involved in transcriptional repression, RNA splicing and ribosome biogenesis. At the protein level most proteins do not appear to display SUMO paralogue-specific modification, while at the site level there is clear evidence of SUMO paralogue specificity. This site-specific selectivity appears, at least in part, to be influenced by proximal amino acids, with generally acidic domains being preferentially modified by SUMO2.

### Experimental procedures

Antibodies and inhibitors. Rabbit antibodies TRIM28 (4124S, 4123S), CTCF (3418S), OCT4A (2890S), SOX2 (23064S), NANOG (3580S), KLF4 (12173S) and mouse antibodies against TRA-1-60 (4746T), TRA-1-81(4745T), SSEA-4 (4755T) and SMA (D4K9N) were from Cell Signalling Technology. The anti-SALL4 (ab29112), anti-TRIM24 (ab70560), anti-NOP58 (ab155556), anti-TRIM24 (ab70560), anti-NESTIN (ab196908) were from Abcam. Mouse antibody against α-Tubulin was from Bethyl Laboratories, mouse anti-LaminA/C antibody was from Sigma (SAB4200236), rabbit anti-mCherry (PA5-34974), rabbit anti-TRIM33 (PA5-82152) and mouse anti-HIS (34650) were from Invitrogen and Qiagen respectively. Anti-CYTOKERATIN17 was a gift from R. Hickerson (University of Dundee). Sheep antibodies against SUMO1, SUMO2 (27) and chicken antibodies against PML (28) were generated in-house. Secondary antibodies conjugated with HRP and Alexa fluorophores were from Sigma and Invitrogen, respectively. ML792 (1644342-14-2), MG132 (474787) and N-ethylmaleimide (E3876) were from Sigma Aldrich. Protease Inhibitor cocktail (11836170001) was from Roche. Propidium iodide and DAPI were from Life Technologies.

### Cloning

SUMO1-KGG-mCherry and SUMO2-KGG-mCherry PiggyBac expression vectors were generated by GATEWAY cloning. Briefly, SUMO1, SUMO2 and mCherry fragments were PCR amplified using the following resources: 6His SUMO1 T95K (300nt) from pSCAI88 and 6His SUMO2 T90K (300nt) from pSCAI89 with a common forwards primer (5’-CACCatgcatcatcatcatcatcatgct-3’) and set of specific mCherry fusing primers (5’-TCACCATACCCCCCTTTTGTTCCTG-3’ and 5’-TCACCATACCTCCCTTCTGCTGCT-3’); mCherry from pRHAI4 CMV-OsTIR1-mCherry2-PURO (700 nt) with a set of common overlapping oligos (5’-GGTATGGTGAGCAAGGGCG-3’ and 5’-TTATTACTTGTACAGCTCGTCCATG-3’).

Subsequently, PCR fragments were fused together using overlap extension PCR and TOPO cloned into pENTR™/D-TOPO™ (Invitrogen) and verified by DNA sequencing. The assembled SUMO1-KGG-mCherry and SUMO2-KGG-mCherry sequences were then sub-cloned from the pENTR vector into the destination PiggyBac GATEWAY expression vector paPX1 (gift from A. Dady UUniversity of Dundee)) using LR clonase (ThermoFisher Scientific).

### Human Induced pluripotent stem cells (hiPSCs) culture and transfection protocols

Human ESC lines (SA121 and SA181) were obtained from Cellartis / Takara Bio Europe. All work with hESCs was approved by the UK Stem cell bank steering committee (Approval reference: SCSC17-14). Human iPSC lines were obtained from Cellartis / Takara Bio Europe (ChiPS4) or the HipSci consortium (bubh3, oaqd3, ueah1 and wibj2). Cell lines were maintained in TESR medium (29) containing FGF2 (Peprotech, 30 ng/ml) and noggin (Peprotech, 10^2^ ng/ml) on growth factor reduced geltrex basement membrane extract (Life Technologies, 10 μg/cm) coated dishes at 37 C in a humidified atmosphere of 5% CO_2_ in air. Cells were routinely passaged twice a week as single cells using TrypLE select (Life Technologies) and replated in TESR medium that was further supplemented with the Rho kinase inhibitor Y27632 (Tocris, 10 μM). Twenty four hours after replating Y27632 was removed from the culture medium. To make SUMO1-KGG-mCherry and SUMO2-KGG-mCherry expressing stable cell lines, ChiPS4 cells were transfected using a Neon electroporation system (Thermo Fisher Scientific) with 10 μl tips. Briefly, ChiPS4 cells were dispersed to single cells as described above then 1×10 cells were collected by centrifugation at 300xg for 2 minutes and resuspended in 11 μl of electroporation buffer R containing 1 μg of either paPX1-SUMO1-KGG-mCherry or paPX1-SUMO2-KGG-mCherry PiggyBac expression vectors along with 0.2 μg of Super PiggyBac transposase (System Biosciences). Electroporation was performed at 1150 V, 1 pulse, 30 mSec and cells plated in mTESR containing Y27632. 5 days after electroporation, mCherry positive cells were positively selected by fluorescence activated cell sorting (FACS) using an SH800 cell sorter (Sony). Monoclonal cell lines were prepared from the bulk sorted population by plating at low density on geltrex coated dishes and individual clones picked using 3.2 mm cloning discs (Sigma Aldrich) soaked in TrypLE select. Cell lines were then expanded and analysed to check for expression of mCherry and His-SUMO1/2.

### Flow cytometry for cell cycle assessment and pluripotency markers

For cell cycle analysis and staining for pluripotency markers ChiPS4 cells were harvested using standard procedures, washed and fixed with ice cold 70% ethanol or 4% PFA for the analysis of cell cycle or NANOG staining respectively. Next cells were stained with propidium iodide or anti-NANOG primary antibody, followed by Alexa 488 conjugated secondary antibody and analysed by flow cytometry using Canto analyser (Becton Dickson). Data was then analysed using FlowJo 10.

### Immunofluorescence, Cell Painting assay and high content microscopy

For IF assays ChiPS4 cells were seeded on µ-Slide 8 Well (Ibidi) or 96 well plates suitable for High Content Microscopy (Nunc). Standard IF procedure was used where appropriate. Briefly, following treatments cells were washed with PBS, fixed with 4% formaldehyde, blocked in 5% BSA in PBS-T and incubated with primary and Alexa conjugated secondary antibodies. Cell Painting was performed as descibed by Bray et al. (30). Imaging and subsequent analysis was performed using IN Cell Analyzer systems (GE Healthcare) and Spotfire (Tibco).

### Protein sample preparation and Western blotting (WB)

ChiPS4 were maintained in a stable culture as described before and treated with inhibitors for a stated time and dose, usually 400nM ML792 was used for 24h or 48h. For WB cells were washed with PBS +/+ and directly lysed in an appropriate volume of 2x Laemmli buffer (approximately 200 μl of buffer was used per 0.5×10^6 cells) (LD; [4% SDS; 20% Glycerol; 120mM 1 M Tris-Cl (pH 6.8); 0.02% w/v bromophenol blue]) and subsequently sonicated using Bioruptor Twin (Diagenode). Protein content was assessed using BCA Protein Assay (ThermoFisher Scientific) and for most purposes 15 μg of total protein was loaded per lane on SDS-Page gel (NuPage 4-12% polyacrylamide, Bis-Tris with MOPS buffer). Proteins were transferred to PVDF membrane using iBlot™ 2 Gel Transfer Device (Invitrogen). Membranes were blocked for 1h in 5% milk in TBS-T and incubated overnight with primary antibodies and 1h with secondary HRP conjugated antibodies before being developed using enhanced chemiluminescence (ThermoFisher Scientific) and exposed to film.

### NiNTA purification

Cells were washed with PBS and scraped in PBS containing 1mM N-ethylmaleimide. The cells were then collected by centrifugation at 300 xg for 5 minutes and the pellets weighed. An aliquot of the cells was lysed in 1.2x NuPage sample buffer (ThermoFisher Scientific) for analysis by Western blotting. The remaining cell pellets (approximately 2 g) were lysed with 5x the pellet weight of lysis buffer (6 M guanidine-HCl, 100 mM sodium phosphate buffer (pH 8.0), 10 mM Tris-HCl (pH 8.0), 10 mM imidazole and 5 mM 2-mercaptoethanol). DNA was sheared by sonication using a probe sonicator (3min, 35% amplitude, 20sec pulses, 20sec intervals on ice and the samples centrifuged at 4000 rpm for 15min at 4 C to remove insoluble material). The protein concentration of the lysate was determined using BCA assay and 6.5mg of total protein from each sample was then incubated overnight at 4°C with 50 μl of packed pre-equlibrated Ni-NTA agarose beads.

After the overnight incubation the supernatant was removed and the beads were washed once with 10 resin volumes of lysis buffer, followed by 1 wash with 10 resin volumes of 8 M urea, 100 mM sodium phosphate buffer (pH 8.0), 10 mM Tris-HCl (pH 8.0), 10 mM imidazole and 5 mM 2-mercaptoethanol, and then 6 washes with 10 resin volumes of 8 M urea, 100 mM sodium phosphate buffer (pH 6.3), 10 mM Tris-HCl (pH 8.0), 10 mM imidazole and 5 mM 2-mercaptoethanol. Proteins were eluted from Ni-NTA agarose beads with 125 μl 1.2x NuPAGE sample buffer for SDS-PAGE.

### Mass Spectrometry based proteomics and quantitative data analysis

Three proteomic experiments are described in this study;

1. Changes in total proteome of ChiPS4 cells during ML792 treatment ChiPS4 cells were either DMSO treated (0 hours condition), or treated with 400nM ML792 for 24 hours or 48 hours. Four replicates of each condition were prepared. Crude cell extracts were made to a protein concentration of between 1 and 2 mg/ml by addition of 1.2x NuPAGE sample buffer to PBS washed cells followed by sonication. For each replicate 25 g protein was fractionated by SDS-PAGE (NuPage 10% polyacrylamide, Bis-Tris with MOPS buffer— Invitrogen) and stained with Coomassie blue. Each lane was excised into four roughly equally sized slices and peptides were extracted by tryptic digestion (31) including alkylation with chloroacetamide. Peptides were resuspended in 35 μL 0.1% TFA 0.5% acetic acid and 10uL of each analysed by LC-MS/MS. This was performed using a Q Exactive mass spectrometer (Thermo Scientific) coupled to an EASY-nLC 1000 liquid chromatography system (Thermo Scientific), using an EASY-Spray ion source (Thermo Scientific) running a 75 μm x 500 mm EASY-Spray column at 45°C. A 240 minute elution gradient with a top 10 data-dependent method was applied. Full scan spectra (m/z 300– 1800) were acquired with resolution R = 70,000 at m/z 200 (after accumulation to a target value of 1,000,000 ions with maximum injection time of 20 ms). The 10 most intense ions were fragmented by HCD and measured with a resolution of R = 17,500 at m/z 200 (target value of 500,000 ions and maximum injection time of 60 ms) and intensity threshold of 2.1×10. Peptide match was set to ‘preferred’, a 40 second dynamic exclusion list was applied and ions were ignored if they had unassigned charge state 1, 8 or >8. Data analysis used MaxQuant version 1.6.1.0 (32). Default setting were used except the match between runs option was enabled, which matched identified peaks among slices from the same position in the gel as well as one slice higher or lower. The uniport human proteome database (downloaded 24/02/2015 - 73920 entries) digested with Trypsin/P was used as search space. LFQ intensities were required for each slice but LFQ normalization was switched off. Manual LFQ normalization was done by calculating the relative LFQ intensity compared to average LFQ intensity for each protein found in the same slice across all 12 samples. For each peptide sample this gave a list of sample LFQ/average LFQ values from which the median was used to normalize all protein LFQ intensities in that sample. The final protein LFQ intensity per lane (and therefore protein sample) was calculated by the sum of normalized LFQ values for that protein intensity in all four slices. Downstream data processing used Perseus v1.6.1.1 (33). Proteins were only carried forward if an LFQ intensity was reported in all four replicates of at least one condition. Zero intensity values were replaced from log2 transformed data (default settings) and outliers were defined by 5% FDR from Student’s t-test using an S0 value of 0.1. A summary of these data can be found in Supp. Data. 1. The mass spectrometry proteomics data have been deposited to the ProteomeXchange Consortium via the PRIDE (34) partner repository with the dataset identifier PXD025867.
2. Characterisation of ChiPS4 cells stably expressing 6His-SUMO1-KGG-mCherry and 6His-SUMO2-KGG-mCherry. Crude cell extracts were prepared in triplicate from parental ChiPS4 cells, ChiPS4-6His-SUMO1-KGG-mCherry and ChiPS4-SUMO2-KGG-mCherry types and fractionated by SDS-PAGE as described above. Samples were prepared and analysed almost identically to experiment 1; gels were sectioned into four slices per lane, tryptic peptides prepared, peptides analysed by LC-MS/MS, and the resultant raw data processed by MaxQuant. The only exceptions were the inclusion of a second sequence database containing the two 6His-SUMO-KGG-mCherry constructs, and the use of MaxQuant LFQ normalization. Two MaxQuant runs were performed; the first aggregating all slices per lane into a single output (“by lane”), and the second considering each slice separately (“by slice”). The former was used to determine cell-specific changes in protein abundance from the proteinGroups.txt file, and the latter used the peptides.txt file to monitor differences in abundance of SUMO-specific peptides between samples, to infer overexpression levels. For the whole cell protein abundance analysis only proteins with data in all three replicates of at least one condition were carried forward. In Perseus zero intensity values were replaced from log2 transformed data (default settings) and outliers were defined by 5% FDR from Student’s t-test using an S0 value of 0.1. A summary of these data can be found in Supp. Data. 2. The mass spectrometry proteomics data have been deposited to the ProteomeXchange Consortium via the PRIDE (34) partner repository with the dataset identifier PXD023241.
3. Identification of SUMO1 and SUMO2 modified proteins from ChiPS4 cells. Two repeats of this experiment were performed using approximately 0.5×10 cells of ChiPS4-6HisSUMO1-KGG-mCherry and ChiPS4-6HisSUMO2-KGG-mCherry per replicate. Samples were taken at different steps of the protocol to assess different fractions. These were; crude cell extracts, NiNTA column elutions and GlyGly-K immunoprecipitations. The last being the source of SUMO-substrate branched peptides. The whole procedure was carried out as described previously (35). In brief, crude cell lysates were prepared of which approximately 100ug was retained for whole proteome analysis as described for experiments 1 and 2 above. The remaining lysate (∼20 mg protein) was used for NiNTA chromatographic enrichment of 6His-SUMO conjugates. Elutions from the NiNTA columns were digested consecutively with LysC then GluC, of which 7% of each was retained for proteomic analysis and the remainder for GlyGly-K immunoprecipitation. The final enriched fractions of LysC and LysC/GluC GG-K peptides were resuspended in a volume of 20 L for proteomic analysis. Peptides from whole cell extracts were analysed once by LC-MS/MS using the same system and settings as described for experiments 1 and 2 above except a 180 minute gradient was used with a top 12 data dependent method. NiNTA elution peptides were analysed identically except a top 10 data dependent method was employed and maximum MS/MS fill time was increased to 120ms. GG-K immunoprecipitated peptides were analysed twice. Firstly, 4uL was fractionated over a 90 minute gradient and analysed using a top 5 data dependent method with a maximum MS/MS fill time of 200ms. Secondly, 11 L of sample was fractionated over a 150 minute gradient and analysed using a top 3 method with a maximum MS/MS injection time of 500ms. Data from WCE and NiNTA elutions were processed together in MaxQuant using Trypsin/P enzyme specificity (2 missed cleavages) for WCE samples and LysC (3 missed cleavages), or LysC+GluC_D/E (considering cleavage after D or E and 8 missed cleavages) for NiNTA elutions. GlyGly (K) and phospho (STY) modifications were selected. The human database and sequences of the two exogenous 6His-SUMO-KGG-mCherry constructs described above were used as search space. In all cases every raw file was treated as a separate ‘experiment’ in the design template such that protein or peptide intensities in each peptide sample were reported, allowing for manual normalization. Matching between runs was allowed but only for peptide samples from the same cellular fraction (WCE, NiNTA elution or GG-K IP), the same or adjacent gel slice, the same protease and the same LC elution gradient. For example, spectra from adjacent gel slices in the WCE fraction across all lanes were matched, and spectra from all GG-K IPs that were digested by the same enzymes were matched. Normalization followed a similar method as described above where ‘equivalent’ peptide samples (ie. those from the same gel slice or ‘equivalent’ peptide samples) from different replicates were compared with one another. For each protein or peptide common to all equivalent peptide samples the intensity in that sample relative to the average across all equivalent samples was calculated. The median of that relative intensity in each peptide sample was used to normalize all protein or peptide intensities from that sample. The final total protein or peptide intensity per replicate was calculated by the sum of all normalized intensities in samples derived from that replicate. It is important to note that peptide samples derived from SUMO1 and SUMO2 cells were considered equivalent for normalization purposes, which assumes largely similar abundances of proteins or peptides between cell types. Zero intensity values were replaced from log2 transformed data (Perseus default settings) and outliers were defined by 5% FDR from Student’s t-test using an S0 value of 0.1. A summary of these data can be found in Supp. Data. 3.

The mass spectrometry proteomics data have been deposited to the ProteomeXchange Consortium via the PRIDE (34) partner repository with the dataset identifier PXD023257.

### Bioinformatic analysis of the SUMO site proteomics

429 proteins identified with at least one SUMO1 or SUMO2 modification site were uploaded to STRING (36) for network analysis. Only proteins associated by a minimum STRING interaction score of 0.7 (high confidence) were included in the final network. Disconnected nodes were removed. Selected groups of functionally related proteins were resubmitted to STRING create smaller sub-networks. These were visualised in Cytoscape v 3.7.2 (37) allowing the graphical display of numbers of sites identified and total GG-K peptide intensity into the protein networks.

### RNA preparation and real-time quantitative PCR (RT-qPCR)

Total RNA was extracted using RNeasy Mini Kit (Qiagen) and treated with the on-column RNase-Free DNase Set (Qiagen) according to the manufacturer’s instructions. RNA concentration was then measured using NanoDrop and 1μg of total RNA per sample was subsequently used to perform a two-step reverse transcription polymerase chain reaction (RT-PCR) using random hexamers and First Strand cDNA Synthesis Kit (ThermoFisher Scientific). Each qPCR reaction contained PerfeCTa SYBR Green FastMix ROX (Quantabio), forward and revers primer mix (200 nM final concentration) and 6 ng of analysed cDNA and was set up in triplicates in MicroAmp™ Fast Optical 96-Well or 384-Well Reaction Plates with Barcodes (Applied Biosystems™). The sequences of primers used were as follows: NANOG (hNANOG_FOR624 ACAGGTGAAGACCTGGTTCC; hNANOG_REV722 GAGGCCTTCTGCGTCACA), SOX2 (hSOX2_FOR907 TGGACAGTTACGCGCACAT; hSOX2_REV1121 CGAGTAGGACATGCTGTAGGT), OCT4A (hOCT4A_FOR825 CCCACACTGCAGCAGATCA and hOCT4A_REV1064 ACCACACTCGGACCACATCC), KLF4 (hKLF4_FOR1630 GGGCCCAATTACCCATCCTT and hKLF4_REV1706 GGCATGAGCTCTTGGTAATGG), TBP (hTBP_FOR896 TGTGCTCACCCACCAACAAT; hTBP_REV1013 TGCTCTGACTTTAGCACCTGTT). Data were collected using QuantStudio™ 6 Flex Real-Time PCR Instrument and analysed using a corresponding software (Applied Biosystems™). Relative amounts of specifically amplified cDNA were calculated using TBP amplicons as normalizers.

Experimental Design and Statistical Rationale All proteomics analyses used a label-free quantitation method employing either triplicate or quadruplicate cell cultures (biological replicates). Biological replicates were individual cell cultures grown and treated separately from one-another but originally derived from a single cell line culture just prior to the experiment. For example, comparisons between different cell lines took individual large-scale cultures for each cell line and split the cells equally among replicate culture dishes.

Once these had grown to appropriate levels of confluence they were treated as described. Intensity data from multiple mass spectrometry runs of the same peptide sample (technical replicates) were normalized against other samples from the same mass spectrometry run and then quantitative data from identical peptide samples aggregated together by summation prior to statistical analyses. Protein or peptide outliers were determined by two-samples student’s t-tests in pairwise comparisons. Specific FDR and S0 values for filtering are described for each experiment above.

## Results

### Inhibition of SUMO modification leads to decreased expression of pluripotency markers in hiPSCs

The pluripotent state in hESCs and hiPSCs is controlled by a network of transcription factors and other chromatin-associated proteins that determine the chromatin environment of key genes (1). To understand the role of SUMO modification in the maintenance of pluripotency in hiPSCs we used ML792 (26) a potent and selective inhibitor of SUMO Activating Enzyme (SAE). This inhibitor has been reported to block proliferation of cancer cells, particularly those overexpressing Myc (26), but has not been evaluated in hiPSCs. To address the role of SUMO modification in hiPSCs, ChiPS4 cells were treated with ML792 in a series of time-course and dose-response experiments. We determined that 400nM ML792 was the lowest concentration that effectively reduced SUMO modification after 4 hrs treatment with minimal effects on cell viability. We restricted our analyses to ML792 treatment times that did not exceed 48 hrs, such that the early effects of SUMO modification inhibition could be evaluated. Microscopic examination revealed that ML792 treatment caused morphological changes with the ChiPS4 cells becoming larger and flatter (Fig. 1a). The rate of proliferation was unchanged after 24 hrs but was slightly reduced after 48 hrs (Fig. 1b). DNA staining of the cells and analysis by flow cytometry indicated that the cell cycle distribution after 24hrs was unaltered by ML792 treatment but after 48 hrs displayed an increased proportion of cells in G2 phase and cells with increased DNA content suggesting endoreplication (Fig. 1c). Western blot analysis revealed a loss of high molecular weight SUMO1 and SUMO2 conjugates and concomitant increase in free SUMOs in ChiPS4 cells (Fig. 1d). This is likely to be a consequence of the rapid removal of SUMO from modified proteins by SUMO specific proteases (SENPs). Analysis of the protein levels of key pluripotency markers indicated that while the levels of OCT4 and SOX2 were unchanged, inhibition of SUMOylation resulted in a decrease in the protein level of NANOG (Fig. 1d).

**Figure 1.**
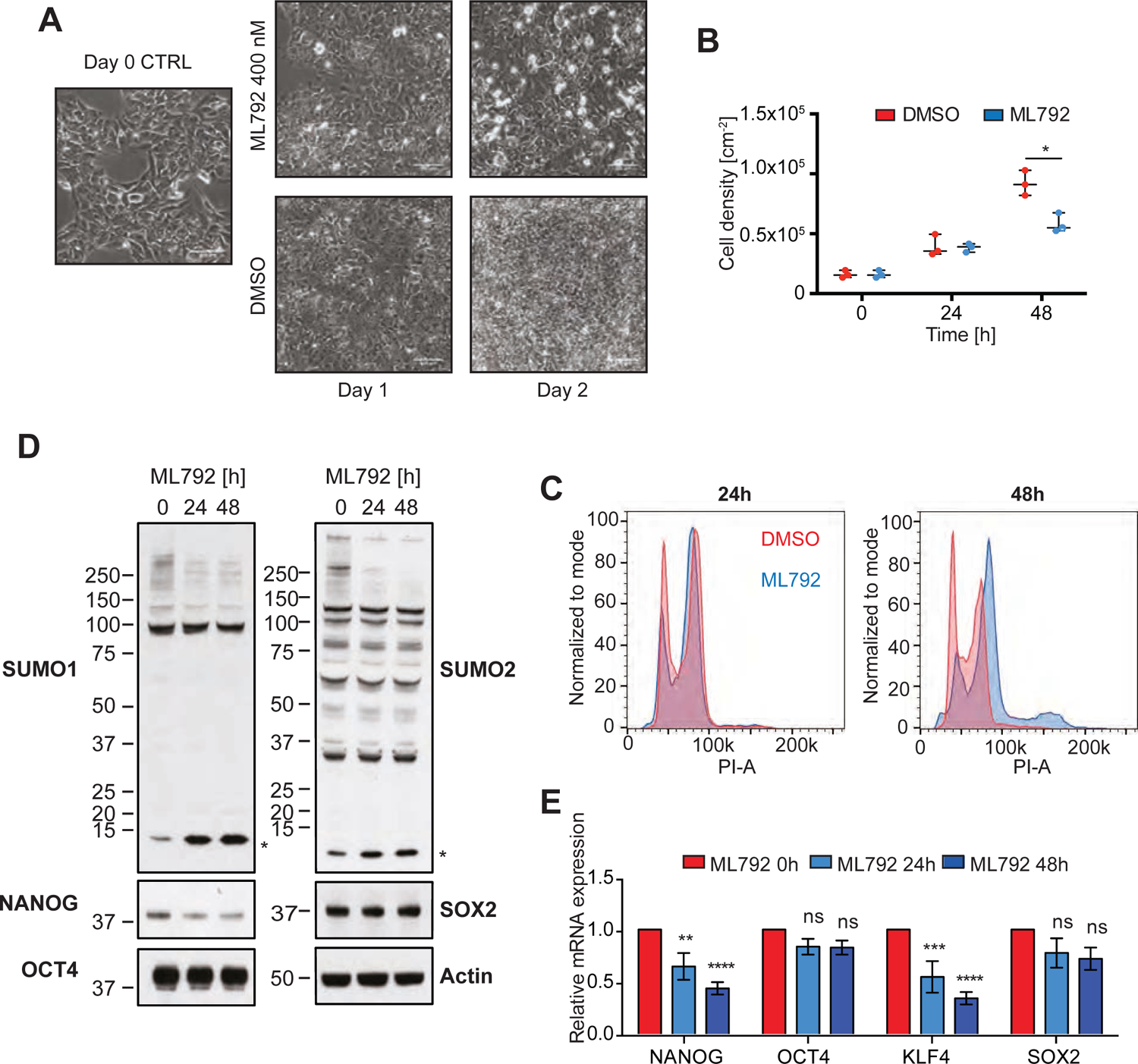
Inhibition of SUMO modification leads to loss of hiPSCs pluripotency. A. ChiPS4 cells were treated with SUMO E1 activating enzyme inhibitor ML792 (400 nM) or DMSO vehicle for the indicated time and analysed for morphology, using phase contrast microscopy (scale bar = 100 μm). B. ChIPS4 cells were seeded at a standard density of 3×10^5^ cells/cm in triplicate and either DMSO or 400nM ML792 treated for the indicated durations. Cells were harvested using TrypleSelect and counted. Data are plotted as a mean (line) ± SEM of individual replicates (dots) n=3. Statistical significance was calculated with t-tests corrected for multiple comparisons using Holm-Sidak’s method (* p<0.05 significantly different from the corresponding DMSO control) C. Crude cell extracts taken from cells incubated at the indicated time-points with 400nM ML792 were analysed by Western blotting to determine conjugation levels of SUMO1, SUMO2/3 and abundance of key pluripotency markers NANOG, SOX2 and OCT4. Anti-Actin western blot was used as a loading control. Asterisk (*) indicates free SUMO1 or SUMO2/3. D. Cells were collected as in B, fixed, stained with propidium iodide (PI) and analysed by flow cytometry. Plots are a representative of three independent experiments. E. Cells were treated with ML792 and after the indicated time total RNA was extracted. mRNA levels of NANOG, OCT4, SOX2 and KLF4 were determined by quantitative PCR. Relative mRNA expression levels normalized to TBP were plotted as means ± SEM of four independent experiments. **p <0.01; *** p <0.001; **** p <0.0001 significantly different from the corresponding value for untreated control (two-way ANOVA followed by Sidak’s multiple comparison test).

This appeared to be a consequence of reduced transcription as determination of mRNA levels by reverse transcriptase quantitative polymerase chain reaction (RT-qPCR) after ML792 treatment demonstrated a reduction in NANOG mRNA. This was also apparent for KLF4, but consistent with the Western blotting results, the levels of OCT4 and SOX2 mRNA were unchanged (Fig. 1e).

### Inhibition of SUMO modification induces phenotypic changes but not large scale proteomic changes in hiPSC

To further investigate the nature and causes of the observed morphological changes induced by the inhibition of SUMO modification, ChiPS4 cells were treated with ML792 for 48h and analysed by phenotypic screening using a cell painting assay (30) (Fig. 2a). Principle component analysis (PCA) indicated that there are clear differences between cells treated with ML792 for 48h and untreated or vehicle (DMSO) treated cells.

**Figure 2.**
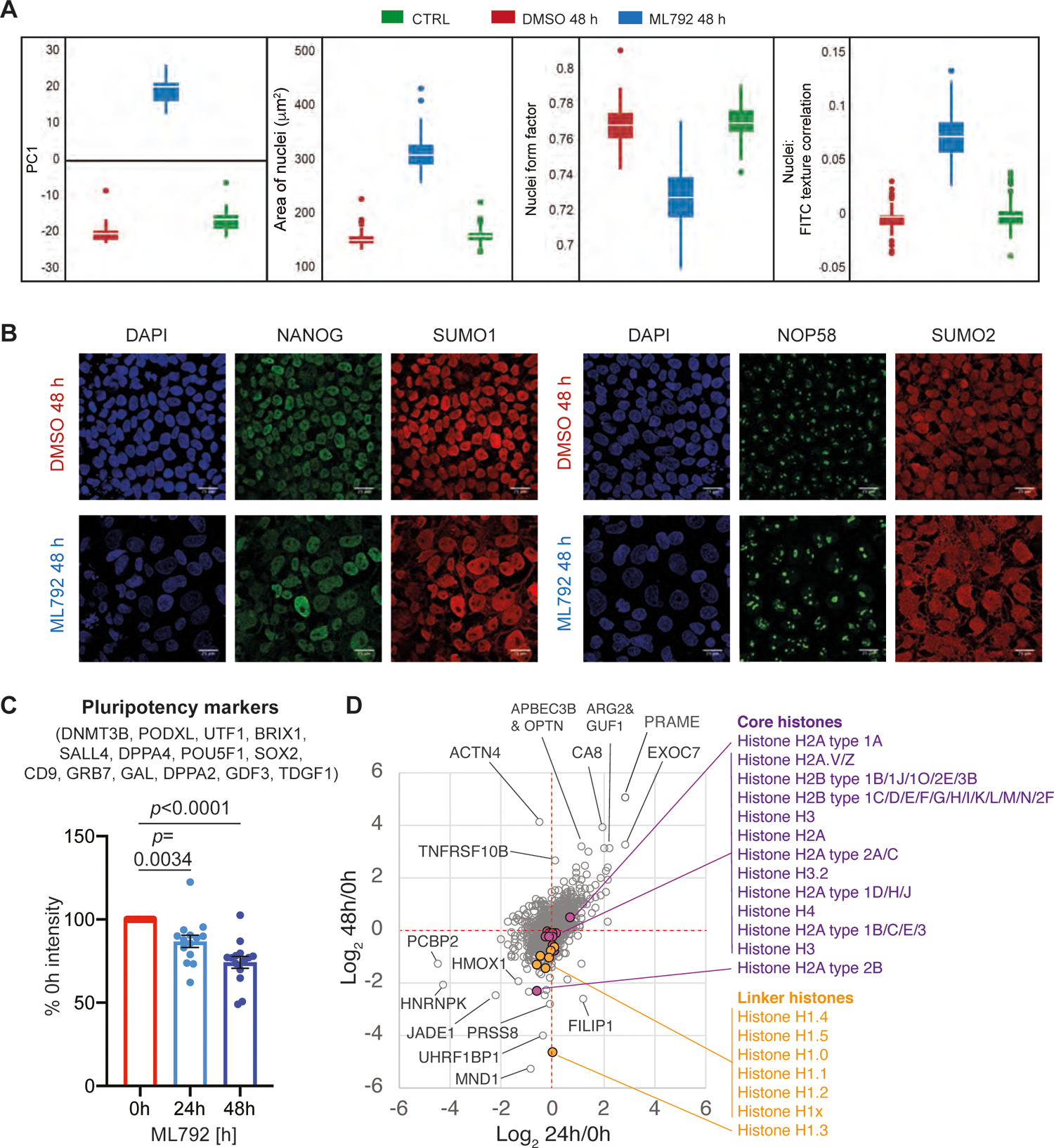
Inhibition of SUMO modification causes morphological changes but does not trigger large-scale proteome changes in hiPSCs. A. Cell painting analysis. ChIPS4 cells were treated with PBS, DMSO vehicle or 400 nM ML792 for 48 h. Cells were then stained, fixed and analysed using high content microscopy. The experiment was performed three times in 8 replicates per condition. Information extracted from Cell Painting analysis was focused on subcellular compartments most affected by ML792 treatment. Most variation between treatments and controls in principal component analysis was captured by PC1. Selected graphs represent quantitation of individual measures contributing to the difference observed in principal component analysis: area of nuclei; nuclei form factor (size and shape of nucleus); nuclei FITC texture correlation (size of nucleolar structures). B. ChIPS4 cells were treated for 48 h with DMSO vehicle or 400 nM ML792, fixed and stained with DAPI (blue), anti-SUMO1 or anti-SUMO2 (red) and anti-NANOG or anti-NOP58 (green) antibodies. Immunofluorescence (IF) images were obtained using Leica SP8 confocal microscope and a 60x water lens. All images contain 25 μm scale bar. C. Crude extracts from hiPSCs treated in triplicate with ML792 for 24h or 48h or not (0h) were analysed by label-free proteomics and the change in protein abundance of detected pluripotency markers (n=14) was calculated as % of 0h intensity. Paired t-test p values are indicated. Average reduction is 13.2% (24h) and 25.7% (48h). D. Scatter plot of Log_2_ 24h/0h and log_2_ 48h/0h abundance change for the entire 4741 protein whole cell proteomic dataset. Extreme outliers are indicated. All identified core and linker histones are represented by coloured markers others are in grey. Linker histones were identified by STRING analysis as a functionally related group of proteins that are significantly reduced in abundance at 48h compared with 0h. Core histone proteins are indicated for reference.

The main differences were found to occur in the nuclear compartment (Fig. 2a). Feature extraction identified changes in the global size (Area of nuclei) and shape (Nuclei form factor) of the nucleus and the structure of the nucleolus (Nuclei: FITC texture correlation) (Fig. 2a). These findings were validated using traditional immunofluorescence (IF) approaches (Fig. 2b). Consistent with previously presented data, NANOG expression as well as the size and shape of the nucleus are both affected by ML792 treatment. NOP58 was used as a marker of the nucleolus, which undergoes a dramatic increase in size and shape (Fig. 2b). As expected, the classic punctate nuclear localisation pattern of SUMO1 and SUMO2 is altered by their deconjugation from substrates, becoming more diffuse and less tightly associated with the nucleus (Fig. 2b).

To evaluate the effect of inhibition of SUMO modification on the global proteome in hiPSCs, total protein extracts were prepared from ChiPS4 cells either untreated or treated with ML792 for 24 or 48 hours and analysed by label-free quantitative proteomics. Data for 4741 proteins was obtained (see Supp. Data File 1). Consistent with SUMO conjugation being critical for pluripotency, known pluripotency markers were modestly but significantly reduced during 24 h and 48 h ML792 exposure (Fig. 2c). However, overall protein abundance changes compared at both time-points showed little evidence for large-scale shifts (Fig. 2d), with few functionally related proteins undergoing coordinated regulation according to STRING: Linker histones appeared to be reduced in abundance after 48 hours, but not 24 hours (Fig. 2d) and proteins from the gene ontology group ‘Collagen-associated extracellular matrix’ had members both up and down-regulated (Supplementary Data file 1). Alone these data do not appear to provide an obvious link between the morphological changes induced by SUMO inhibition and underlying protein abundance changes. This implies that it is an accumulation of multiple small protein abundance changes effected by or in addition to deSUMOylation of key regulators, that explains the consequences of ML792 treatment on hiPSCs.

### Generation of hiPSC lines for SUMO proteomic analysis

Our data suggest an important role for SUMO in maintaining the pluripotent state of hiPSCs. Given that previous studies of SUMOylation have typically focused on somatic cells, it was important to establish the SUMOylome of hiPSCs to identify candidate proteins that either directly or indirectly contribute to the observed phenotype. We adapted a proteomic approach that allows sites in proteins modified by SUMO1 and SUMO2/3 to be identified (38). To enable this analysis in hiPSCs the ChIPS4 cell line was engineered to stably express 6His-SUMO-mCherry constructs for either SUMO1 or SUMO2 (Fig. 3a, b) that incorporated the TGG to KGG mutations to facilitate GlyGly-K peptide immunoprecipitation and identification as described previously (38). As mCherry is linked to the C-terminus of SUMO the expressed fusion protein will be processed by endogenous SUMO proteases to expose the C-terminal GlyGly sequence for conjugation and release free mCherry. Single cell clones were selected based on the expression of free mCherry and the level of SUMO paralogue expression (Fig. 3c). Clones selected for SUMO site proteomic experiments were extensively characterised to ensure that the exogenous SUMO was functional and that the cells retained their pluripotency. Western blotting indicated that His-tagged SUMO-KGG paralogues were conjugated to substrates in response to heat shock (Supplementary Fig. 1). Cells expressing SUMO1-KGG and SUMO2-KGG had normal cell cycle profiles (Supplementary Fig. 2a), expressed levels of pluripotency markers comparable to wild type ChIPS4 cells (Supplementary Fig. 2b, c) and retained the ability to differentiate into endoderm, ectoderm and mesoderm (Supplementary Fig. 2d). Analysis of the whole cell proteomes (Fig. 3d) of ChIPS4 cells expressing SUMO1-KGG and SUMO2-KGG identified the expected exogenous mCherry, SUMO1 and SUMO2 peptides (Fig. 3e) while peptides common to both endogenous and exogenous SUMOs showed both types were conjugated to substrates at roughly similar levels (Fig. 3f). Importantly, the engineered cell lines did not show large-scale differences from parental cells in their expressed proteomes (Fig. 3g, h, see also Supplementary Data File 2). Together these data indicate that expression of SUMO mutants did not disrupt the normal pluripotent state or differentiation potential of ChiPS4 cells.

**Figure 3.**
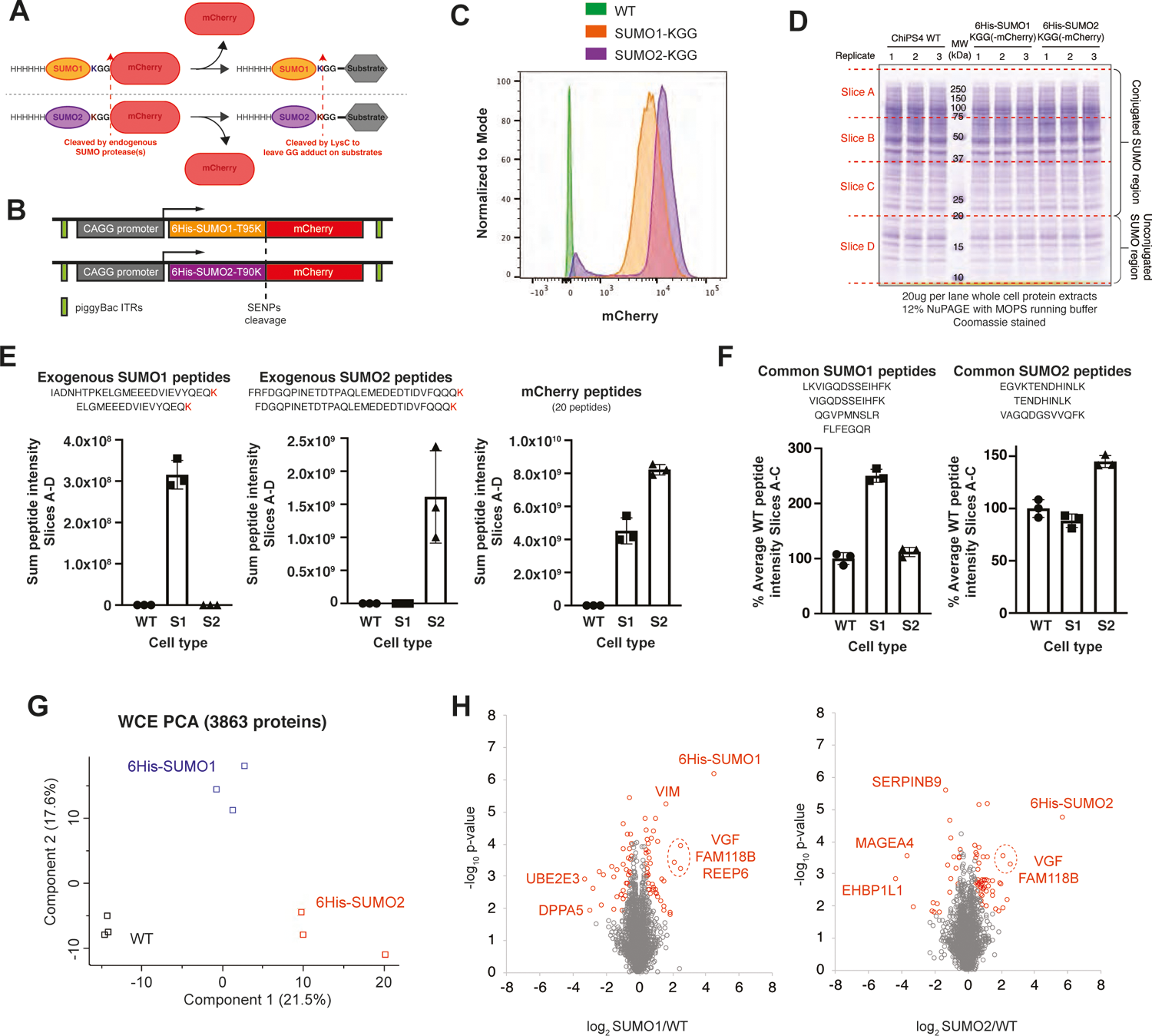
ChiPS4 cell lines engineered for SUMO1 and SUMO2 site proteomic analysis. A. Overview of the principles behind the SUMO-mCherry construct design and the utility of the expressed proteins for SUMO site proteomic analysis in CHiPS4 cells. The C-terminal mCherry protein used for cell selection is cleaved away from 6His-SUMO by endogenous SUMO proteases allowing conjugation to substrate proteins. B. Scheme representing the piggyBac construct design used for generation of ChiPS4 cell lines expressing 6His-SUMO1-KGG-mCherry or 6His-SUMO2-KGG-mCherry. C. Flow cytometry analysis of single cell clones using mCherry expression levels to infer SUMO expression levels. D. Coomassie stained SDS-PAGE gel fractionating whole cell protein extracts from parental CHiPS4 cells (WT) and the selected 6His-SUMO-KGG-mCherry clones prepared in triplicate. This was used to monitor SUMO overexpression levels using proteomic analysis and each lane was excised into 4 sections allowing differentiation between conjugated (slices A-C) and unconjugated (Slice D) SUMO forms. E. Peptide intensity data from slices A-D for two peptides each from 6His-SUMO1-KGG (left) and 6His-SUMO2-KGG (centre) that are unique to the exogenous constructs. Data for 20 mCherry peptides are also shown (right). F. Peptide intensity data from slices A-C for four peptides from SUMO1 (left) and three from SUMO2 (right) that are common to both the endogenous and exogenous forms of the proteins. G. Quantitative data from 3863 proteins identified from the gel shown in D were compared by principal component analysis. H. Numerical ratio and unpaired student’s t-test results comparing WT parental cells with 6His-SUMO1-KGG-mCherry cells (left), and WT with 6His-SUMO2-KGG-mCherry cells (right). Outliers (red markers) were defined in Perseus by 5% FDR with an S0 value of 0.1 (78 outliers from WT vs SUMO1 cells and 64 from WT vs SUMO2 cells). Gene names from extreme outliers are indicated.

### Identification of SUMO1 and SUMO2 targets in hiPSCs

The workflow for the identification of SUMO1 and SUMO2 targets incorporates proteomic analysis at three levels (Supplementary Fig. 3a). This involved analysis of whole cell extracts to monitor total protein levels, analysis of nickel affinity purified proteins to identify SUMO modified proteins and analysis of GG-K immunoprecipitations to define sites of SUMO modification. SUMO1/SUMO2 comparisons can be made at the site level because after LysC digestion both mutants leave the same GG-K adduct on substrates (Supplementary. Fig. 3b), and so intensity comparisons can be used to infer preference for SUMO type at the site level. The experiment was conducted twice in triplicate and PCA indicated that replicates performed at the same time displayed a high degree of clustering (Supplementary Fig. 3c). PCA also indicated clear differences between SUMO1 and SUMO2 as well as between experiments carried out at different times (Supplementary Fig. 3c). This is likely to be a reflection of the precise growth state of the cells when the experiment was conducted. Very few total protein abundance differences between SUMO1 and SUMO2 cell were consistent to both experimental runs, but there were substantial and consistent differences between SUMO1 and SUMO2 at both nickel affinity purifications and at the site level (Supplementary Fig. 3d). Across the two experimental runs and aggregating the data for SUMO1 and SUMO2 a total of 976 SUMO sites were identified in 427 proteins. Approximately 84% of these had already been described in at least one of four large-scale SUMO2 site proteomics studies totalling 49768 unique sites of non-stem cell origin (see Supplementary Data file 3) (Fig. 4a). DNA methyl transferase 3B (DNMT3B) and the key embryonic stem cell transcription factor SALL4 were among a small group of proteins with at least three novel sites in this study (Fig. 4a). Although peptide intensity is a relatively imprecise proxy for abundance, it has been successfully used for SUMO-substrate branched peptides to separate high occupancy from low occupancy sites (39). For our data total GG-K peptide intensity suggests SALL4 is among the most modified SUMO substrate in hiPSCs and contains 17 sites of modification (Fig. 4b). DNMT3B contains 12 sites and is also among the most abundantly modified substrates, while the methyl DNA binding protein MBD1 contains 8 sites and is also in the top cohort of substrates by modified peptide intensity (Fig. 4b). Transcription intermediary factors 1 α and 1 β (TRIM24 and TRIM28) are also both highly modified substrates. The boundary and chromatin isolation factor CTCF is heavily modified with SUMO, consistent with a role for SUMO in chromatin architecture (Fig. 4b). There is also evidence for extensive SUMO chain formation as the branch points from SUMO2/3 chains are amongst the most abundant GG-K peptides (Fig. 4b). Strikingly, the SUMOylated forms of a number of these nuclear substrates could be detected directly in total whole cell lysates from control ChiPS4 cells, but not ML792 treated hiPSCs (Fig. 4c), implicating SUMO as a major regulator of the total pool of these proteins. STRING enrichment analysis of the 427 modified proteins created a network consisting of 3 broad clusters that could be categorised as having functions in ribosome biogenesis, RNA splicing, and regulation of gene expression (Fig. 4d). Despite forming extensive protein networks (Fig. 5a, b), proteins involved in ribosome biogenesis and RNA splicing represented only approximately 5% of the total GGK peptide intensity (Fig. 4b insert). Most of the remainder have roles in transcription and chromatin structure or are closely linked to these functions (Fig. 5c-h). There is a prominent network of zinc-finger transcription factors, closely associated with TRIM28 (Fig. 5c) which contains many of the most heavily SUMOylated proteins identified. The TRIM-ZNF-SUMO axis may play a key role in silencing retroviral elements and this may be particularly important in hiPSCs (40). Histone proteins themselves form a small cluster of SUMO substrates (Fig. 5d) which STRING positioned in the centre of the gene regulation region of the whole network (Fig. 4c). The transcriptional regulators themselves form a bipolar network (Fig. 5e) with the smaller sub-cluster consisting mainly of apparently weakly modified ribosomal proteins and the larger sub-cluster containing many heavily modified chromatin associated proteins. Strikingly, many members of chromatin remodelling complexes such as PRC2 (Fig. 5f), BAF (Fig. 5g) and NURD (Fig. 5h) are among this group. Together these networks and clusters of proteins provide multiple direct and indirect links between SUMO and chromatin structure regulation.

**Figure 4.**
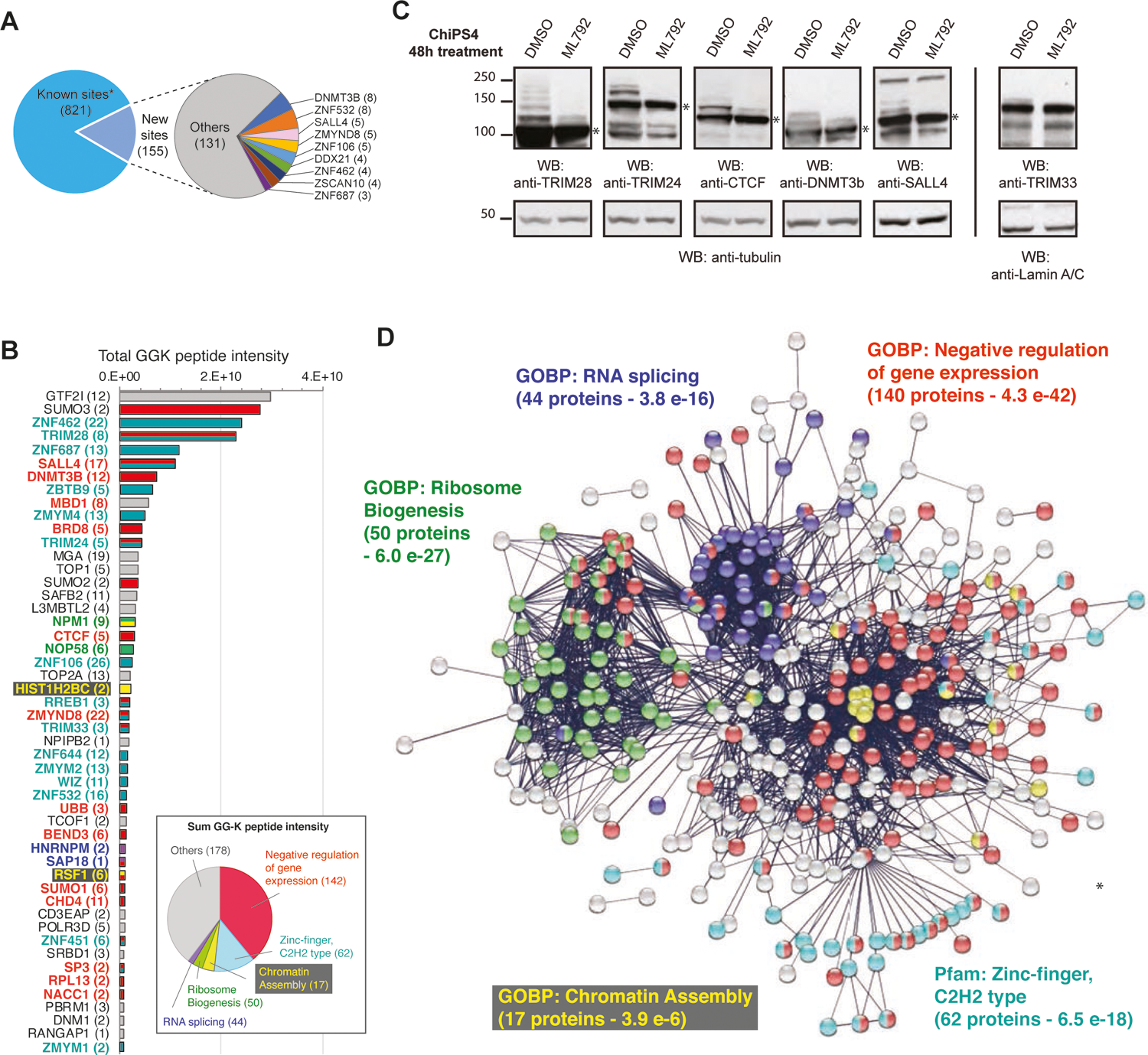
Functions of SUMO1 and SUMO2 targets in hiPSCs. A. 976 SUMO sites were identified from 6His-SUMO1-KGG and 6His-SUMO2-KGG IPS cells, of which 155 were novel compared with previous high-throughput SUMO site proteomics studies (See Supplementary Data File 2). Proteins with three or more novel sites are highlighted. B. Summary of the top 50 SUMO substrates by total SUMO1+SUMO2 GGK-peptide intensity for all identified sites. Gene names are shown with numbers of sites in brackets. Bars are colour coded by category shown in panel C. Insert shows contribution to total GGK peptide intensity of proteins from the categories shown in C (note categories are not mutually exclusive). C. STRING interaction network of the 427 IPS SUMO substrates. Only high confidence interactions were considered from ‘Text mining’, ‘Experiments’ and ‘Databases’ sources. Network PPI enrichment p-value <1.0 ×10. Nodes are coloured by functional or structural group as indicated. D. Immunoblot analysis of ChIPS4 cells treated with the SUMO E1 conjugating enzyme inhibitor ML792 or DMSO for 48h. Total protein extracts were probed with anti-TRIM28, anti-TRIM24, anti-CTCF, anti-DNMT3b, anti-SALL4, anti-TRIM33 and anti-tubulin or anti-Lamin A/C antibodies (loading controls). SUMO-modified species above the band for unmodified proteins (*), reduce with ML792 treatment.

**Figure 5.**
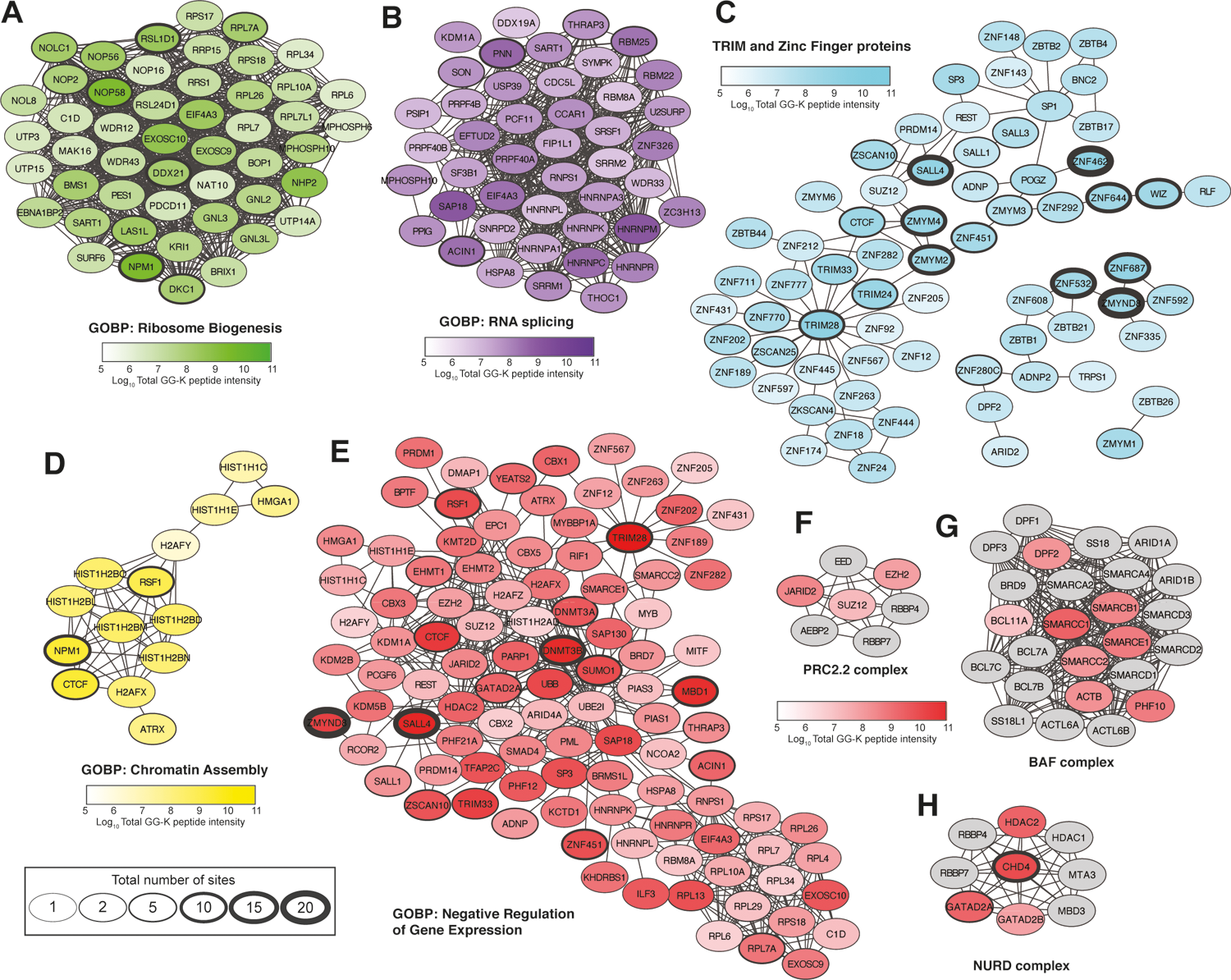
Networks of functionally related SUMO substrates in hiPSCs. A-E. Detailed protein interaction networks derived from the broad categories shown in Fig. 3C. Node shade is proportional to log_10_ total GGK peptide intensity and border thickness indicates numbers of sites found. F-H shows data for selected chromatin remodelling complexes. Grey nodes were not identified in the present study but included to complete complexes. TRIM proteins were included in C to link more zinc-finger proteins in the network.

### Differences between SUMO1 and SUMO2 at the protein and acceptor site level

The proteomic experimental design allowed quantitative comparisons between SUMO1 and SUMO2 at multiple stages of the purification process (Supplementary Fig. 3a): SUMO1/SUMO2 ratios from crude IPS cell extracts shows there to be few differences (0.03% significant) at the whole proteome level (Fig. 6a). There are also surprisingly few differences (7.8% significant) between NiNTA purifications from the two cell types (Fig. 6a). Exceptions include the well-documented SUMO1 substrate RanGAP1, along with TRIM24 and TRIM33 which all appear to show similar levels of SUMO1 preference (Fig. 6a). In contrast, over half of the GGK-containing peptides quantified in both experimental runs and both cell lines showed large and significant difference between SUMO1 and SUMO2 cells (Fig. 6a). Extreme examples of SUMO1 preferential sites include not only RanGAP1 K524, but also TRIM33 K776, NFRKB K359, PNN K157, TFPT K216 and two sites in TOP1 (K117 & K134) (Fig. 6a). Conversely, TRIM28 contains two of the most SUMO2-preferntial sites at K507 and K779, and lysines 48 and 63 from ubiquitin also seems to be among the extreme SUMO2 acceptors (Fig. 6a). In the context of whole proteins, three key proteins in our dataset, SALL4, TRIM24 and TRIM33 show largely SUMO1 preferential modification, while TRIM28 and CTCF show apparent preference for SUMO2 (Fig. 6b, c). This was broadly confirmed by Western-blot analysis from NiNTA purifications (Fig. 6d), although in some cases the degree of preference was not as striking as expected. Significantly, Western-blot analysis of NiNTA purifications confirmed the SUMO2-modified form of CTCF to be more abundant than SUMO1, but both the modified and unmodified forms of CTCF were purified (Fig. 6d). Thus, it seems likely that large amounts of unmodified CTCF obscured the SUMO2 preference in the NiNTA purification proteomic data. This effect may be rare, but highlights potential weaknesses in proteomics studies when using relatively low-stringency purifications such as 6His/Ni-NTA pull downs.

**Figure 6.**
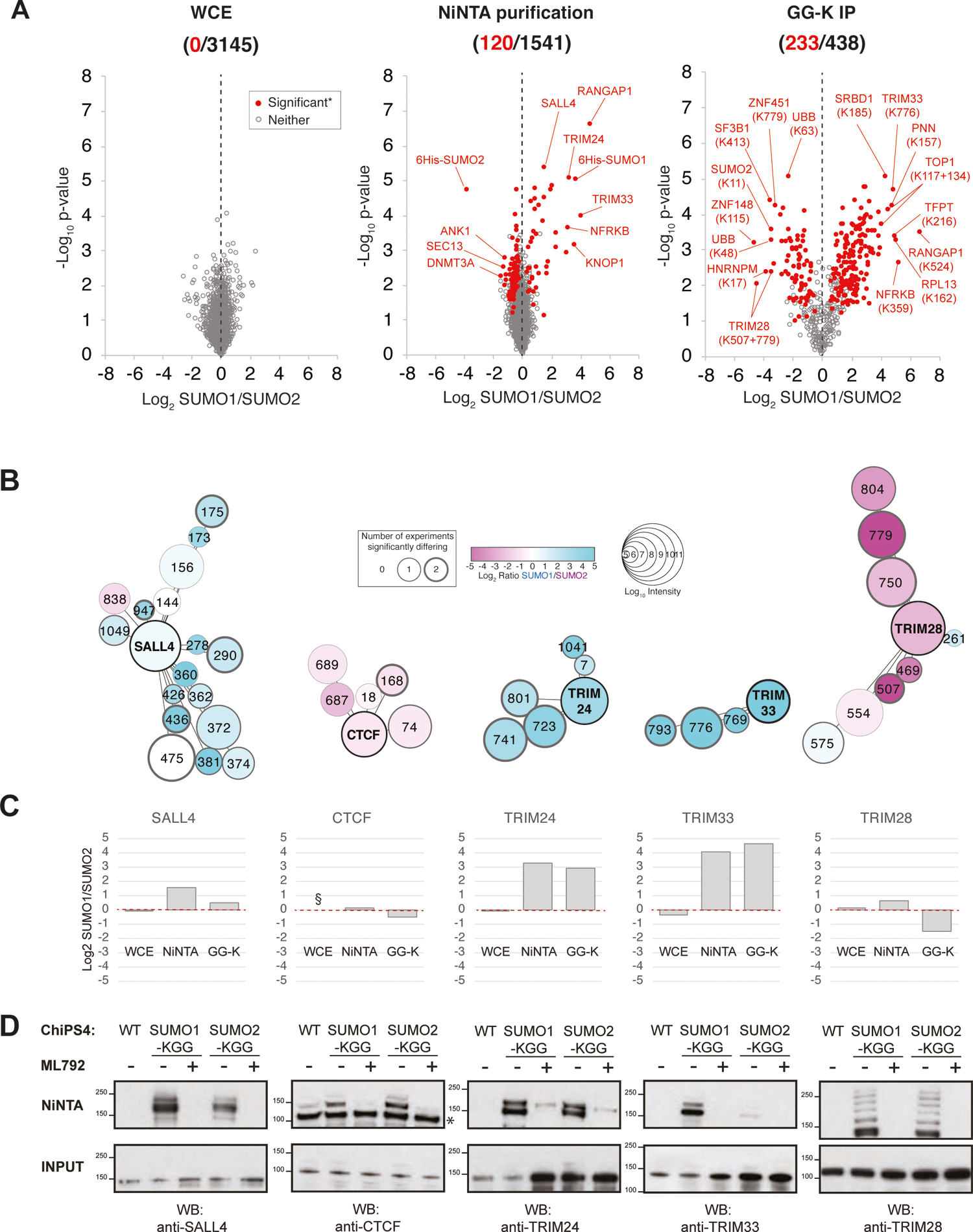
SUMO paralogue preference at the site level in hiPSCs is influenced by proximal acidic residues. A. Scatter plots of log_2_ SUMO1/SUMO2 ratio and -log_10_ student’s t-test p-value for proteins or peptides detected in the different cellular fractions. Red markers were found to be significantly different in both experimental runs. Selected outliers are indicated. Numbers of significantly differing proteins or peptides compared to the entire set of proteins or peptides quantified are shown. B. Schematic presentations of selected proteins found to have high total GG-K peptide intensities in the present study. Site positions are indicated by number. SUMO preference is represented by colour and GGK peptide intensity by the size of the site node (see key). Site node border line thickness represents number of experiments in which it was found to show a significant SUMO preference. Protein nodes are labelled with the gene name and are colour is based on SUMO preference calculated by total GGK peptide intensity from SUMO1 cells/total from SUMO2 cells. Edges linking sites to proteins are angled relative to their position in the linear protein sequence with first and last residues at the top of the protein node. C. Summary of log2 SUMO1/SUMO2 intensity measured in each of the three cellular fractions analysed. § - No data acquired. D. Immunoblot analysis of whole cell extracts (input) and NiNTA purifications from SUMO1 and SUMO2 cells either treated or not with 400nM ML792. * - Position of unmodified protein in NiNTA purifications.

### Preference for SUMO2 modification in regions of higher negative charge

The proteomic data reveal many multiply modified proteins to have a strong overall SUMO paralog selection. TOP1, TRIM33, ZNF462 and ZNF532 all contain multiple SUMO1-specific sites, and conversely, DNMT3a, CHD4, ubiquitin and SUMOs 2 and 3 themselves have a majority of SUMO2-specific sites (Supplementary Fig. 4). However, a number of substrates such as GTF2I, SAFB2 and MGA (Supplementary Fig. 4) have a mixture of sites showing varied SUMO type preference. This broad range of site-specific SUMO paralogue preference raises the question of how specificity is determined. By averaging the SUMO1/SUMO2 peptide intensity ratio data over the two experimental runs we ranked 739 sites by paralogue preference from SUMO1 to SUMO2 (Fig. 7a). Not all sites could be included for sequence analysis as sites close to protein N or C termini lacked amino acids in their sequence windows. Sequence logos of the top and bottom 123 sites (one sixth of total) shows broadly similar motifs for SUMO1 and SUMO2 sites (Fig. 7b), with both showing the characteristic ψKxE motif. However, outside the consensus SUMO2 sites are more prone to D or E in the −2 position than SUMO1 sites, and a significant number of SUMO1 sites have P in +3 (Fig. 7b). More generally, D/E residues are more enriched within the 21 residue sequence windows of SUMO2 than SUMO1. This is confirmed by sequence window charge analysis (Fig. 7c) which shows acceptor lysine residues preferentially modified with SUMO2 over SUMO1 are more likely to be embedded in a sequence that is net negatively charged at pH7.4 than those showing SUMO1 preference. This trend is also generally progressive across the spectrum of site-specific SUMO paralogue selection (Supplementary. Fig. 5a-e). For some proteins where SUMO paralogue preference varies in different regions of the sequence, this coincides with local charge variations consistent with SUMO2 preference in acidic domains. SALL4, CHD4 and SAFB (Fig. 7d) show that sites preferentially modified by SUMO2 were generally negatively charged while SUMO1-preferential modification took place in regions of the protein of that were more positively charged.

**Figure 7.**
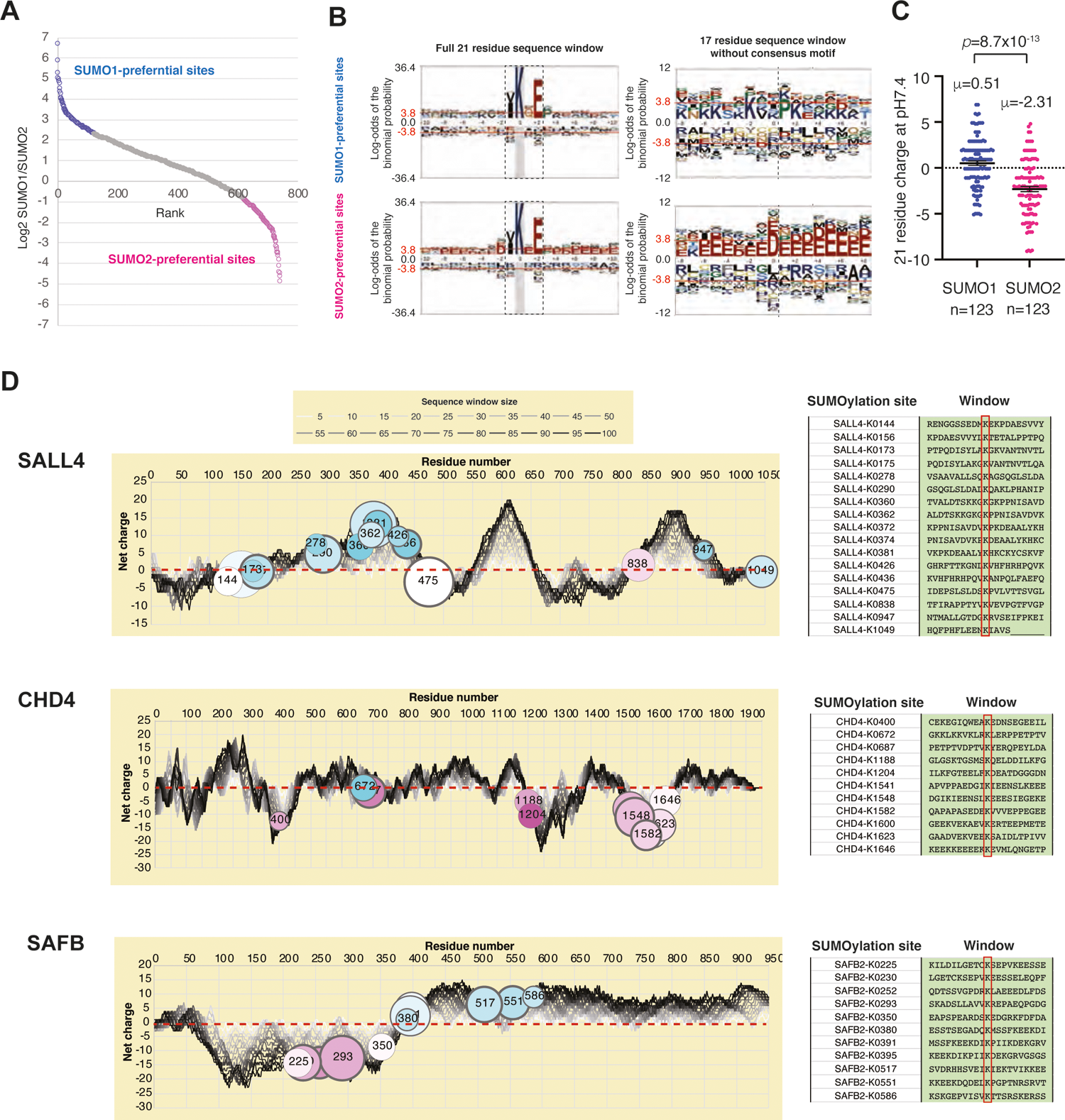
Validation of SUMOylation status of abundant SUMO substrates in ChIPS4 cells. A. Log_2_ SUMO1/SUMO2 ratio versus rank of ratio for 738 GG-K containing peptides that could be numerically compared between SUMO1 and SUMO2 samples. Top and bottom (123 sites) by ratio are selected to represent SUMO paralogue-preferential sites (coloured) with the remainder marked in grey. B. pLogos (52) for the 123 SUMO1 or SUMO2 specific sites as shown in part C using the human proteome as background. Two analyses were performed using the entire 21 residue window (left), and the 17 residue window of the same sequences without the 4 residue SUMO consensus motif (right). The position of the consensus sequence region is boxed with a dashed line (left) or shown by a dotted line (right). C. Comparison between SUMO1 and SUMO2-preferential sites for predicted net charge at pH7.4 of the amino-acid region encompassing −10 to +10 residues around the acceptor lysine. Charge predicted using Isoelectric Point Calculator tool (53). D. Schematic presentations summarising site-specific SUMOylation data, and predicted sequence charge distribution for SALL4, CHD4 and SAFB2. Net charge at each amino acid position in the sequence (left) was calculated for sequence windows of sizes from 5 to 100 residues as indicated in the key (see M&M for details). SUMO conjugation sites and 21 residue sequence windows are shown (right).

## Discussion

Our studies highlight an important role for SUMO in maintaining the pluripotent state of ChIPS4 hiPSCs. Using a potent and highly specific inhibitor of the E1 SUMO Activating Enzyme ML792 (26) we blocked de novo SUMO modification which allowed endogenous SUMO specific proteases to remove SUMO from previously modified proteins. In this way ML792 treatment effectively deSUMOylated cellular proteins. In response to 48h of 400nM ML792 exposure ChIPS4 cells showed reduced pluripotency and underwent dramatic morphological changes, particularly to nuclear structure and size. Over the same period of time there appeared to be few large-scale changes to the cellular proteome, implying that the consequences of loss of SUMOylation in hiPSCs primarily involves functional changes to factors critical for nuclear structure and function, probably followed by multiple modest protein abundance changes, which together lead to the observed phenotype. Candidates for these SUMOylated factors were identified by SUMO site proteomic analysis. The major network that accounted for the bulk of SUMO modification sites was associated with negative regulation of gene expression. At the heart of this network is TRIM28, the histone methyl transferase SETDB1 and the Chromobox silencer CBX3. ChIP-seq derived locations of TRIM28, SETDB1 and CBX3 indicate that they are associated with retroviral elements (41). These proteins along with SUMO have previously been shown to function in HERV silencing in mouse ES cells (40, 42) and adult human cells (43, 44) and this is consistent with our proteomic analysis that indicates that all three of these proteins are heavily SUMO modified (Fig. 4). The TRIM28 co-repressor functions by interacting with DNA bound Kruppel type Zinc finger proteins and these proteins are also identified as being SUMO modified in our proteomic studies and are part of the large TRIM28 centric network of SUMO modified proteins. It is also suggested that a number of developmental genes are repressed by TRIM28/KRAB-ZNFs through H3K9me3 and de novo DNA methylation of their promoter regions (45), thus making TRIM28/ZNFs a crucial link in maintenance of pluripotency in human stem cells. These data are consistent with the wide distribution of SUMO on chromatin (46–49) and with a very different distribution in murine fibroblasts and embryonic stem cells (9). Recently Theurillat et al.(50) reported a SUMO site proteomic analysis of mouse embryonic stem cells. In this case the method used allowed analysis of endogenous SUMO2 but not SUMO1 modification sites. The number of sites identified in this study was rather similar (608 SUMO2 sites in 350 proteins) to that reported here (976 SUMO1 plus SUMO2 sites in 427 proteins), as were the major functional networks. However, the precise targets identified were very different, with only SALL4 of the top 20 mouse ES cell specific proteins being present in the top 50 (based on peptide intensity) SUMO modified proteins in human stem cells reported here (Fig. 4b). Such differences presumably reflect the differences between mESCs and hiPSCs and the developmental stages which these cells represent. While the mouse cells were established from blastocysts the human cells are derived by reprogramming normal somatic cells (25).

Analysis of the SUMO proteome of the ChIPS4 hiPSCs shows that well-defined groups of protein are modified. Aside from the TRIM28/ZNF network mentioned above, proteins involved in “ribosome biogenesis” and “splicing” are SUMO modified and this likely impacts on the normal growth and self-renewal of the hiPSCs. However, the largest network of proteins falls into the category of “negative regulation of transcription”, with many chromatin remodellers, chromatin modification and DNA modification enzymes identified as SUMO substrates. Thus, SUMO modification is likely to play an important role in maintaining pluripotency of hiPSCs by repressing genes that either disrupt pluripotency or drive differentiation.

During our analysis we also found that a large proportion of modification sites displayed a clear preference for either SUMO1 or SUMO2, and that this seems to be influenced by the charge distribution around the acceptor lysine. Specifically, lysines within acidic domains are more likely to be modified by SUMO2 than SUMO1. While this is by no means a strict rule, these data suggest site-level SUMO paralog selection is at least in part defined by the electrostatic environment of the acceptor lysine. Intriguingly, structures of Ubc9 in complex with either SUMO1 or SUMO2 (51) show significant charge deviation between the two paralogues close to the active site of the E2 enzyme (Supplementary Fig. 6), potentially implicating the SUMO molecule itself in site-selection. It is important to note however, that in spite of the clear variety of site-level SUMO paralogue preference, most proteins display little overall SUMO preference when considering the total cellular pool together, meaning examples of entirely SUMO1 or SUMO2/3 modified proteins are probably rare. Understanding the differences between SUMO1 and SUMO2/3 in terms of site-selectivity and downstream functional outcomes will likely be critical in gaining a full understanding of the role of SUMOylation in all higher eukaryotic cell types.

## Abbreviations

bFGF: basic fibroblast growth factor

ESCs: embryonic stem cells

FACS: fluorescence activated cell sorting

GGK: GlyGly-Lys branched peptide

hESCs: human embryonic stem cells

hiPSCs: human induced pluripotent stem cells

LIF: leukemia inhibitory factor

mESCs: mouse embryonic stem cells

## Acknowledgements

We would like to acknowledge the invaluable help from various facilities at the University of Dundee: Flow Cytometry and Cell Sorting, National Phenotypic Screening Centre, Dundee Imaging Facility, Human Pluripotent Stem Cell Facility. This work was supported by an Investigator Award from Wellcome (217196/Z/19/Z) and a Programme grant from Cancer Research UK (C434/A21747) to RTH. ML is part of the Ubi-Code European Training Network that received funding from the European Union Horizon 2020 research and innovation programme under the Marie Curie grant agreement No. 765445. BM was supported by the European Union’s Horizon 2020 research and innovation programme under the Marie Skłodowska-Curie grant agreement No. 704989.

## Author Contributions

B.M. cloned expression vectors, generated cell lines, designed and performed most experiments using hiPSCs and interpreted data. ML performed NiNTA purifications and Western blotting analyses. L.D. was in charge of cell culture, cell line derivation and quality control and was consulted over experimental design. M.H.T consulted over proteomic experimental design, acquired and processed MS data and conducted bioinformatic and statistical analyses. E.B. performed bioinformatic analysis of the structural data. B.M., M.H.T., L.D., and R.T.H. contributed to data analysis. B.M., R.T.H., M.H.T. and L.D. wrote the paper. R.T.H. conceived the project.

### Data availability

Data underlying all Figures and Supplementary Figures are available in the source data file. Source data for proteomics experiments are deposited in PRIDE as described in the Methods section. All other data are available from the corresponding authors on reasonable request.

### Supplementary data files

This article contains supplemental data including following supplementary data files:

Supplementary data file 1. Summary of the quantitative data from the proteomics experiment to study changes to the cellular proteome during ML792 treatment of ChiPS4 cells.

Supplementary data file 2. Summary of the quantitative data from the proteomics experiment to study differences in the cellular proteome among wild type ChiPS4 cells and cells expressing 6His-SUMO1-KGG-mCherry or 6His-SUMO2-KGG-mCherry.

Supplementary data file 3. Summary of the quantitative data from the proteomics experiment to identify SUMO1 and SUMO2 targets from ChiPS4 cells.

## Author Information

Correspondence and requests for materials should be addressed to R.T.H

## Notes

### Competing Interest Statement

The authors have declared no competing interest.

### Summary of Updates

There is a significant change in the manuscript due to adaptations we had to make to fit with the demands of editors. Due to the fact that the focus of the manuscript has shifted completely to proteomics data, we had to amend as much as the authors list and title of the paper.

## Bibliography

1. Young, R. A. (2011) Control of the embryonic stem cell state. Cell 144, 940–954

2. Takahashi, K., Tanabe, K., Ohnuki, M., Narita, M., Ichisaka, T., Tomoda, K., and Yamanaka, S. (2007) Induction of pluripotent stem cells from adult human fibroblasts by defined factors. Cell 131, 861–872

3. Takahashi, K., and Yamanaka, S. (2006) Induction of pluripotent stem cells from mouse embryonic and adult fibroblast cultures by defined factors. Cell 126, 663–676

4. Yu, J., Vodyanik, M. A., Smuga-Otto, K., Antosiewicz-Bourget, J., Frane, J. L., Tian, S., Nie, J., Jonsdottir, G. A., Ruotti, V., Stewart, R., Slukvin, II, and Thomson, J. A. (2007) Induced pluripotent stem cell lines derived from human somatic cells. Science 318, 1917–1920

5. Apostolou, E., and Hochedlinger, K. (2013) Chromatin dynamics during cellular reprogramming. Nature 502, 462–471

6. Vierbuchen, T., and Wernig, M. (2012) Molecular roadblocks for cellular reprogramming. Mol Cell 47, 827–838

7. Borkent, M., Bennett, B. D., Lackford, B., Bar-Nur, O., Brumbaugh, J., Wang, L., Du, Y., Fargo, D. C., Apostolou, E., Cheloufi, S., Maherali, N., Elledge, S. J., Hu, G., and Hochedlinger, K. (2016) A Serial shRNA Screen for Roadblocks to Reprogramming Identifies the Protein Modifier SUMO2. Stem Cell Reports 6, 704–716

8. Cheloufi, S., Elling, U., Hopfgartner, B., Jung, Y. L., Murn, J., Ninova, M., Hubmann, M., Badeaux, A. I., Euong Ang, C., Tenen, D., Wesche, D. J., Abazova, N., Hogue, M., Tasdemir, N., Brumbaugh, J., Rathert, P., Jude, J., Ferrari, F., Blanco, A., Fellner, M., Wenzel, D., Zinner, M., Vidal, S. E., Bell, O., Stadtfeld, M., Chang, H. Y., Almouzni, G., Lowe, S. W., Rinn, J., Wernig, M., Aravin, A., Shi, Y., Park, P. J., Penninger, J. M., Zuber, J., and Hochedlinger, K. (2015) The histone chaperone CAF-1 safeguards somatic cell identity. Nature 528, 218–224

9. Cossec, J. C., Theurillat, I., Chica, C., Bua Aguin, S., Gaume, X., Andrieux, A., Iturbide, A., Jouvion, G., Li, H., Bossis, G., Seeler, J. S., Torres-Padilla, M. E., and Dejean, A. (2018) SUMO Safeguards Somatic and Pluripotent Cell Identities by Enforcing Distinct Chromatin States. Cell Stem Cell 23, 742–757 e748

10. Hendriks, I. A., and Vertegaal, A. C. (2016) A comprehensive compilation of SUMO proteomics. Nat Rev Mol Cell Biol 17, 581–595

11. Mahajan, R., Delphin, C., Guan, T., Gerace, L., and Melchior, F. (1997) A small ubiquitin-related polypeptide involved in targeting RanGAP1 to nuclear pore complex protein RanBP2. Cell 88, 97–107

12. Hay, R. T. (2005) SUMO: a history of modification. Mol Cell 18, 1–12

13. Flotho, A., and Melchior, F. (2013) Sumoylation: a regulatory protein modification in health and disease. Annu Rev Biochem 82, 357–385

14. Rodriguez, M. S., Dargemont, C., and Hay, R. T. (2001) SUMO-1 conjugation in vivo requires both a consensus modification motif and nuclear targeting. J Biol Chem 276, 12654–12659

15. Sampson, D. A., Wang, M., and Matunis, M. J. (2001) The small ubiquitin-like modifier-1 (SUMO-1) consensus sequence mediates Ubc9 binding and is essential for SUMO-1 modification. J Biol Chem 276, 21664–21669

16. Tatham, M. H., Jaffray, E., Vaughan, O. A., Desterro, J. M., Botting, C. H., Naismith, J. H., and Hay, R. T. (2001) Polymeric chains of SUMO-2 and SUMO-3 are conjugated to protein substrates by SAE1/SAE2 and Ubc9. J Biol Chem 276, 35368–35374

17. Song, J., Durrin, L. K., Wilkinson, T. A., Krontiris, T. G., and Chen, Y. (2004) Identification of a SUMO-binding motif that recognizes SUMO-modified proteins. Proc Natl Acad Sci U S A 101, 14373–14378

18. Williams, R. L., Hilton, D. J., Pease, S., Willson, T. A., Stewart, C. L., Gearing, D. P., Wagner, E. F., Metcalf, D., Nicola, N. A., and Gough, N. M. (1988) Myeloid leukaemia inhibitory factor maintains the developmental potential of embryonic stem cells. Nature 336, 684–687

19. Ying, Q. L., Nichols, J., Chambers, I., and Smith, A. (2003) BMP induction of Id proteins suppresses differentiation and sustains embryonic stem cell self-renewal in collaboration with STAT3. Cell 115, 281–292

20. Humphrey, R. K., Beattie, G. M., Lopez, A. D., Bucay, N., King, C. C., Firpo, M. T., Rose-John, S., and Hayek, A. (2004) Maintenance of pluripotency in human embryonic stem cells is STAT3 independent. Stem Cells 22, 522–530

21. Xu, C., Rosler, E., Jiang, J., Lebkowski, J. S., Gold, J. D., O’Sullivan, C., Delavan-Boorsma, K., Mok, M., Bronstein, A., and Carpenter, M. K. (2005) Basic fibroblast growth factor supports undifferentiated human embryonic stem cell growth without conditioned medium. Stem Cells 23, 315–323

22. Xu, R. H., Chen, X., Li, D. S., Li, R., Addicks, G. C., Glennon, C., Zwaka, T. P., and Thomson, J. A. (2002) BMP4 initiates human embryonic stem cell differentiation to trophoblast. Nat Biotechnol 20, 1261–1264

23. James, D., Levine, A. J., Besser, D., and Hemmati-Brivanlou, A. (2005) TGFbeta/activin/nodal signaling is necessary for the maintenance of pluripotency in human embryonic stem cells. Development 132, 1273–1282

24. Xu, R. H., Peck, R. M., Li, D. S., Feng, X., Ludwig, T., and Thomson, J. A. (2005) Basic FGF and suppression of BMP signaling sustain undifferentiated proliferation of human ES cells. Nat Methods 2, 185–190

25. Rossant, J. (2015) Mouse and human blastocyst-derived stem cells: vive les differences. Development 142, 9–12

26. He, X., Riceberg, J., Soucy, T., Koenig, E., Minissale, J., Gallery, M., Bernard, H., Yang, X., Liao, H., Rabino, C., Shah, P., Xega, K., Yan, Z.-h., Sintchak, M., Bradley, J., Xu, H., Duffey, M., England, D., Mizutani, H., Hu, Z., Guo, J., Chau, R., Dick, L. R., Brownell, J. E., Newcomb, J., Langston, S., Lightcap, E. S., Bence, N., and Pulukuri, S. M. (2017) Probing the roles of SUMOylation in cancer cell biology by using a selective SAE inhibitor. Nature Chemical Biology 13, 1164–1171

27. Tatham, M. H., Geoffroy, M. C., Shen, L., Plechanovova, A., Hattersley, N., Jaffray, E. G., Palvimo, J. J., and Hay, R. T. (2008) RNF4 is a poly-SUMO-specific E3 ubiquitin ligase required for arsenic-induced PML degradation. Nat Cell Biol 10, 538–546

28. Hands, K. J., Cuchet-Lourenco, D., Everett, R. D., and Hay, R. T. (2014) PML isoforms in response to arsenic: high-resolution analysis of PML body structure and degradation. J Cell Sci 127, 365–375

29. Ludwig, T. E., Levenstein, M. E., Jones, J. M., Berggren, W. T., Mitchen, E. R., Frane, J. L., Crandall, L. J., Daigh, C. A., Conard, K. R., Piekarczyk, M. S., Llanas, R. A., and Thomson, J. A. (2006) Derivation of human embryonic stem cells in defined conditions. Nat Biotechnol 24, 185–187

30. Bray, M. A., Singh, S., Han, H., Davis, C. T., Borgeson, B., Hartland, C., Kost-Alimova, M., Gustafsdottir, S. M., Gibson, C. C., and Carpenter, A. E. (2016) Cell Painting, a high-content image-based assay for morphological profiling using multiplexed fluorescent dyes. Nat Protoc 11, 1757–1774

31. Shevchenko, A., Tomas, H., Havlis, J., Olsen, J. V., and Mann, M. (2006) In-gel digestion for mass spectrometric characterization of proteins and proteomes. Nat Protoc 1, 2856–2860

32. Cox, J., and Mann, M. (2008) MaxQuant enables high peptide identification rates, individualized p.p.b.-range mass accuracies and proteome-wide protein quantification. Nat Biotechnol 26, 1367–1372

33. Tyanova, S., Temu, T., Sinitcyn, P., Carlson, A., Hein, M. Y., Geiger, T., Mann, M., and Cox, J. (2016) The Perseus computational platform for comprehensive analysis of (prote)omics data. Nat Methods 13, 731–740

34. Perez-Riverol, Y., Csordas, A., Bai, J., Bernal-Llinares, M., Hewapathirana, S., Kundu, D. J., Inuganti, A., Griss, J., Mayer, G., Eisenacher, M., Perez, E., Uszkoreit, J., Pfeuffer, J., Sachsenberg, T., Yilmaz, S., Tiwary, S., Cox, J., Audain, E., Walzer, M., Jarnuczak, A. F., Ternent, T., Brazma, A., and Vizcaino, J. A. (2019) The PRIDE database and related tools and resources in 2019: improving support for quantification data. Nucleic Acids Res 47, D442–D450

35. Tammsalu, T., Matic, I., Jaffray, E. G., Ibrahim, A. F., Tatham, M. H., and Hay, R. T. (2015) Proteome-wide identification of SUMO modification sites by mass spectrometry. Nat Protoc 10, 1374–1388

36. Szklarczyk, D., Gable, A. L., Lyon, D., Junge, A., Wyder, S., Huerta-Cepas, J., Simonovic, M., Doncheva, N. T., Morris, J. H., Bork, P., Jensen, L. J., and Mering, C. V. (2019) STRING v11: protein-protein association networks with increased coverage, supporting functional discovery in genome-wide experimental datasets. Nucleic Acids Res 47, D607–D613

37. Shannon, P., Markiel, A., Ozier, O., Baliga, N. S., Wang, J. T., Ramage, D., Amin, N., Schwikowski, B., and Ideker, T. (2003) Cytoscape: a software environment for integrated models of biomolecular interaction networks. Genome Res 13, 2498–2504

38. Tammsalu, T., Matic, I., Jaffray, E. G., Ibrahim, A. F. M., Tatham, M. H., and Hay, R. T. (2014) Proteome-wide identification of SUMO2 modification sites. Sci Signal 7, rs2

39. Hendriks, I. A., Lyon, D., Young, C., Jensen, L. J., Vertegaal, A. C., and Nielsen, M. L. (2017) Site-specific mapping of the human SUMO proteome reveals co-modification with phosphorylation. Nat Struct Mol Biol 24, 325–336

40. Wolf, D., and Goff, S. P. (2007) TRIM28 Mediates Primer Binding Site-Targeted Silencing of Murine Leukemia Virus in Embryonic Cells. Cell 131, 46–57

41. Gerstein, M. B., Kundaje, A., Hariharan, M., Landt, S. G., Yan, K. K., Cheng, C., Mu, X. J., Khurana, E., Rozowsky, J., Alexander, R., Min, R., Alves, P., Abyzov, A., Addleman, N., Bhardwaj, N., Boyle, A. P., Cayting, P., Charos, A., Chen, D. Z., Cheng, Y., Clarke, D., Eastman, C., Euskirchen, G., Frietze, S., Fu, Y., Gertz, J., Grubert, F., Harmanci, A., Jain, P., Kasowski, M., Lacroute, P., Leng, J. J., Lian, J., Monahan, H., O’Geen, H., Ouyang, Z., Partridge, E. C., Patacsil, D., Pauli, F., Raha, D., Ramirez, L., Reddy, T. E., Reed, B., Shi, M., Slifer, T., Wang, J., Wu, L., Yang, X., Yip, K. Y., Zilberman-Schapira, G., Batzoglou, S., Sidow, A., Farnham, P. J., Myers, R. M., Weissman, S. M., and Snyder, M. (2012) Architecture of the human regulatory network derived from ENCODE data. Nature 489, 91–100

42. Yang, B. X., El Farran, C. A., Guo, H. C., Yu, T., Fang, H. T., Wang, H. F., Schlesinger, S., Seah, Y. F., Goh, G. Y., Neo, S. P., Li, Y., Lorincz, M. C., Tergaonkar, V., Lim, T. M., Chen, L., Gunaratne, J., Collins, J. J., Goff, S. P., Daley, G. Q., Li, H., Bard, F. A., and Loh, Y. H. (2015) Systematic identification of factors for provirus silencing in embryonic stem cells. Cell 163, 230–245

43. Schmidt, N., Domingues, P., Golebiowski, F., Patzina, C., Tatham, M. H., Hay, R. T., and Hale, B. G. (2019) An influenza virus-triggered SUMO switch orchestrates co-opted endogenous retroviruses to stimulate host antiviral immunity. Proceedings of the National Academy of Sciences 116, 17399–17408

44. Tie, C. H., Fernandes, L., Conde, L., Robbez-Masson, L., Sumner, R. P., Peacock, T., Rodriguez-Plata, M. T., Mickute, G., Gifford, R., Towers, G. J., Herrero, J., and Rowe, H. M. (2018) KAP1 regulates endogenous retroviruses in adult human cells and contributes to innate immune control. EMBO Rep 19

45. Oleksiewicz, U., Gladych, M., Raman, A. T., Heyn, H., Mereu, E., Chlebanowska, P., Andrzejewska, A., Sozanska, B., Samant, N., Fak, K., Auguscik, P., Kosinski, M., Wroblewska, J. P., Tomczak, K., Kulcenty, K., Ploski, R., Biecek, P., Esteller, M., Shah, P. K., Rai, K., and Wiznerowicz, M. (2017) TRIM28 and Interacting KRAB-ZNFs Control Self-Renewal of Human Pluripotent Stem Cells through Epigenetic Repression of Pro-differentiation Genes. Stem Cell Reports 9, 2065–2080

46. Neyret-Kahn, H., Benhamed, M., Ye, T., Le Gras, S., Cossec, J. C., Lapaquette, P., Bischof, O., Ouspenskaia, M., Dasso, M., Seeler, J., Davidson, I., and Dejean, A. (2013) Sumoylation at chromatin governs coordinated repression of a transcriptional program essential for cell growth and proliferation. Genome Res 23, 1563–1579

47. Niskanen, E. A., Malinen, M., Sutinen, P., Toropainen, S., Paakinaho, V., Vihervaara, A., Joutsen, J., Kaikkonen, M. U., Sistonen, L., and Palvimo, J. J. (2015) Global SUMOylation on active chromatin is an acute heat stress response restricting transcription. Genome Biol 16, 153

48. Seifert, A., Schofield, P., Barton, G. J., and Hay, R. T. (2015) Proteotoxic stress reprograms the chromatin landscape of SUMO modification. Sci Signal 8, rs7

49. Liu, H. W., Zhang, J., Heine, G. F., Arora, M., Gulcin Ozer, H., Onti-Srinivasan, R., Huang, K., and Parvin, J. D. (2012) Chromatin modification by SUMO-1 stimulates the promoters of translation machinery genes. Nucleic Acids Res 40, 10172–10186

50. Theurillat, I., Hendriks, I. A., Cossec, J. C., Andrieux, A., Nielsen, M. L., and Dejean, A. (2020) Extensive SUMO Modification of Repressive Chromatin Factors Distinguishes Pluripotent from Somatic Cells. Cell Rep 32, 108146

51. Gareau, J. R., Reverter, D., and Lima, C. D. (2012) Determinants of small ubiquitin-like modifier 1 (SUMO1) protein specificity, E3 ligase, and SUMO-RanGAP1 binding activities of nucleoporin RanBP2. J Biol Chem 287, 4740-4751

52. O’Shea, J. P., Chou, M. F., Quader, S. A., Ryan, J. K., Church, G. M., and Schwartz, D. (2013) pLogo: a probabilistic approach to visualizing sequence motifs. Nat Methods 10, 1211–1212

53. Kozlowski, L. P. (2016) IPC - Isoelectric Point Calculator. Biol Direct 11, 55

