## Supplementary Figures and Legends for "Identification of SUMO targets required to maintain human stem cells in the pluripotent state"

**Supplementary Figures and Figure Legends S1-S6**

**Supplementary data files**

This article contains supplemental data that are included as separate data files that contain following information:

***Supplementary data file 1.*** Summary of the quantitative data from the proteomics experiment to study changes to the cellular proteome during ML792 treatment of ChiPS4 cells.

***Supplementary data file 2.*** Summary of the quantitative data from the proteomics experiment to study differences in the cellular proteome among wild type ChiPS4 cells and cells expressing 6His-SUMO1-KGG-mCherry or 6His-SUMO2-KGG-mCherry.

***Supplementary data file 3.*** Summary of the quantitative data from the proteomics experiment to identify SUMO1 and SUMO2 targets from ChiPS4 cells.

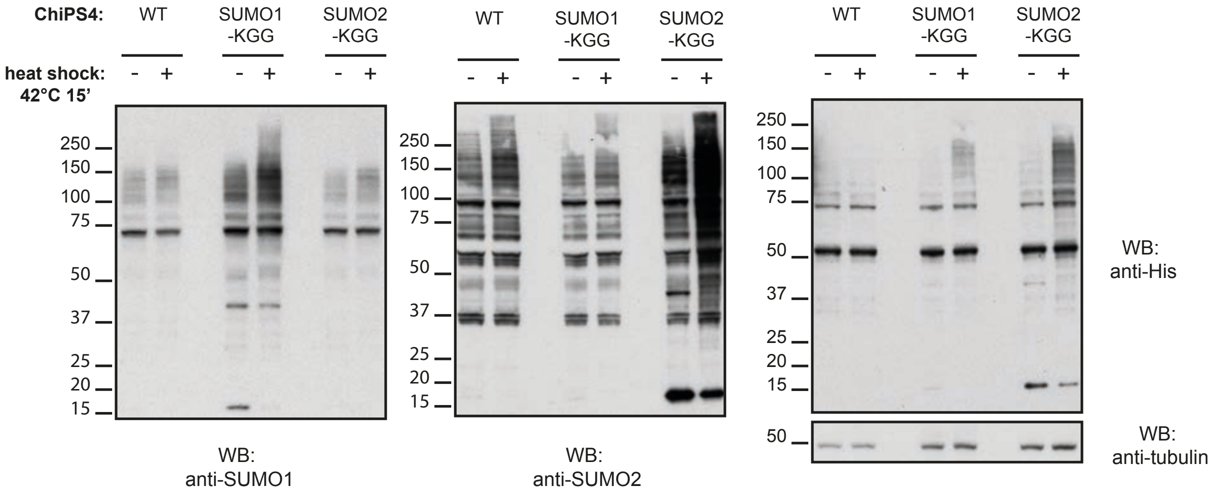


**Supplementary Figure 1. Exogenous 6His-SUMOKGG constructs do not significantly affect pluripotency or the cellular proteome of hiPSCs.**

ChiPS4 WT, SUMO1-KGG and SUMO2-KGG expressing cell lines were exposed to heat shock for 15 minutes at 42°C and total protein lysates were analysed by western blot using anti-SUMO1, anti-SUMO2/3, anti-His and anti-tubulin (loading control) antibodies.

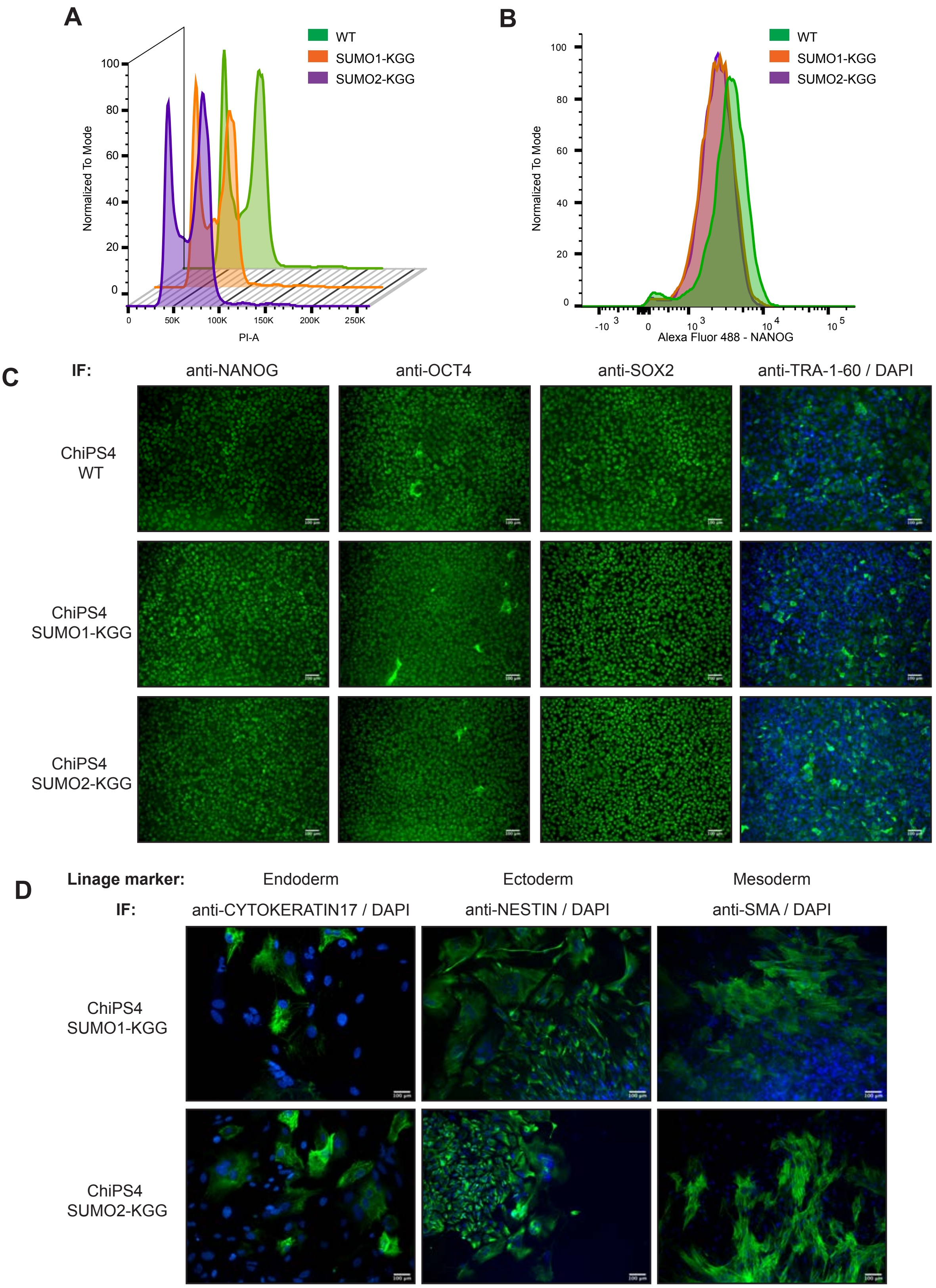


**Supplementary Figure 2. hiPSCs expressing 6His-SUMOKGG constructs do not show any cell cycle, pluripotency or differentiation defects.**

Flow cytometry analysis of **A.** cell cycle and **B.** NANOG expression. **C.** Immunofluorescence analysis of pluripotency associated markers (NANOG, SOX2, OCT4, TRA-1-60) in ChiPS4 WT, SUMO1-KGG and SUMO2-KGG expressing cell lines. **D.** *In vitro* differentiation potential of ChiPS4 SUMO1-KGG and SUMO2-KGG expressing cell lines was assessed by immunofluorescence staining with DAPI and specific antibodies against CYTOKERATIN 17 (Endoderm), NESTIN (Ectoderm) and SMA (Mesoderm). **C – D.** Immunofluorescence (IF) images were obtained using a Leica DM-IRB microscope equipped with a Hamamatsu CCD camera and 20x 0.3C-Plan lens. All images contain 100 μm scale bar.

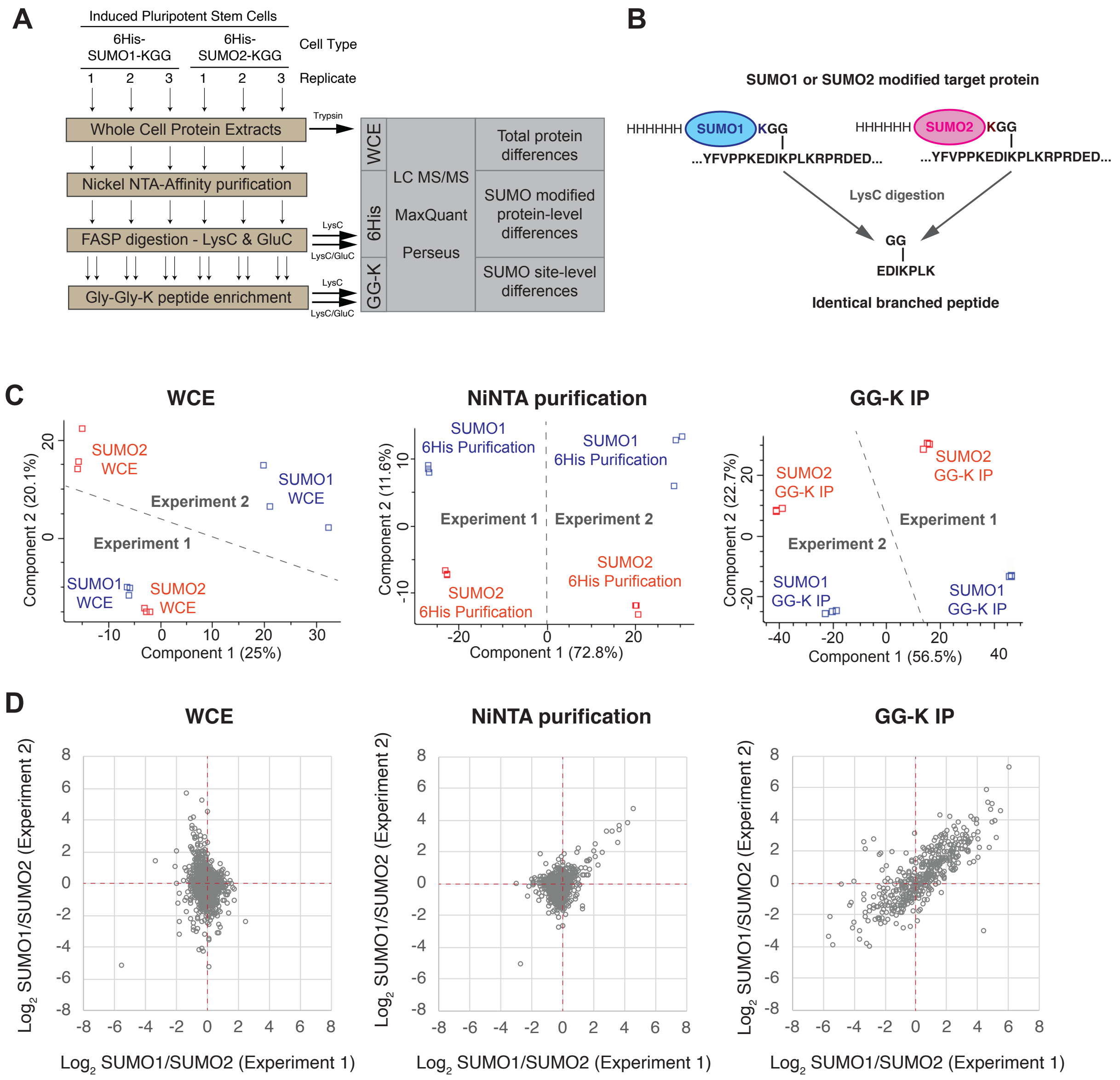


**Supplementary Figure 3. Overview of experimental design and proteomic data relating to SUMO1 and SUMO2 site identification in hiPSCs**

**A.** Overview of a proteomics experiment to identify IPS-specific SUMO1 and SUMO2 substrates. Two experimental runs were performed with two different hiPSC lines (expressing 6His-SUMO1-KGG or 6His-SUMO2-KGG), each one was performed in triplicate. Three protein fractions were analysed; whole cell extracts (WCE), NiNTA column elutions (6HIS), GlyGly-K immunoprecipitated peptide elutions (GG-K IP). All peptides were analysed by LC-MS/MS and data processed by MaxQuant. **B.** SUMO1-KGG and SUMO2-KGG proteins leave identical GG adducts on substrates after LysC digestion, therefore peptide intensity differences between cell types can be used to infer site-specific SUMO paralogue preference. **C.** Principal component analyses of MS data from the three different cell fractions. **D.** Comparisons between experimental runs for SUMO1/SUMO2 ratio data for each of the three cell fractions analysed.

SUMO targets in human induced pluripotent stem cells

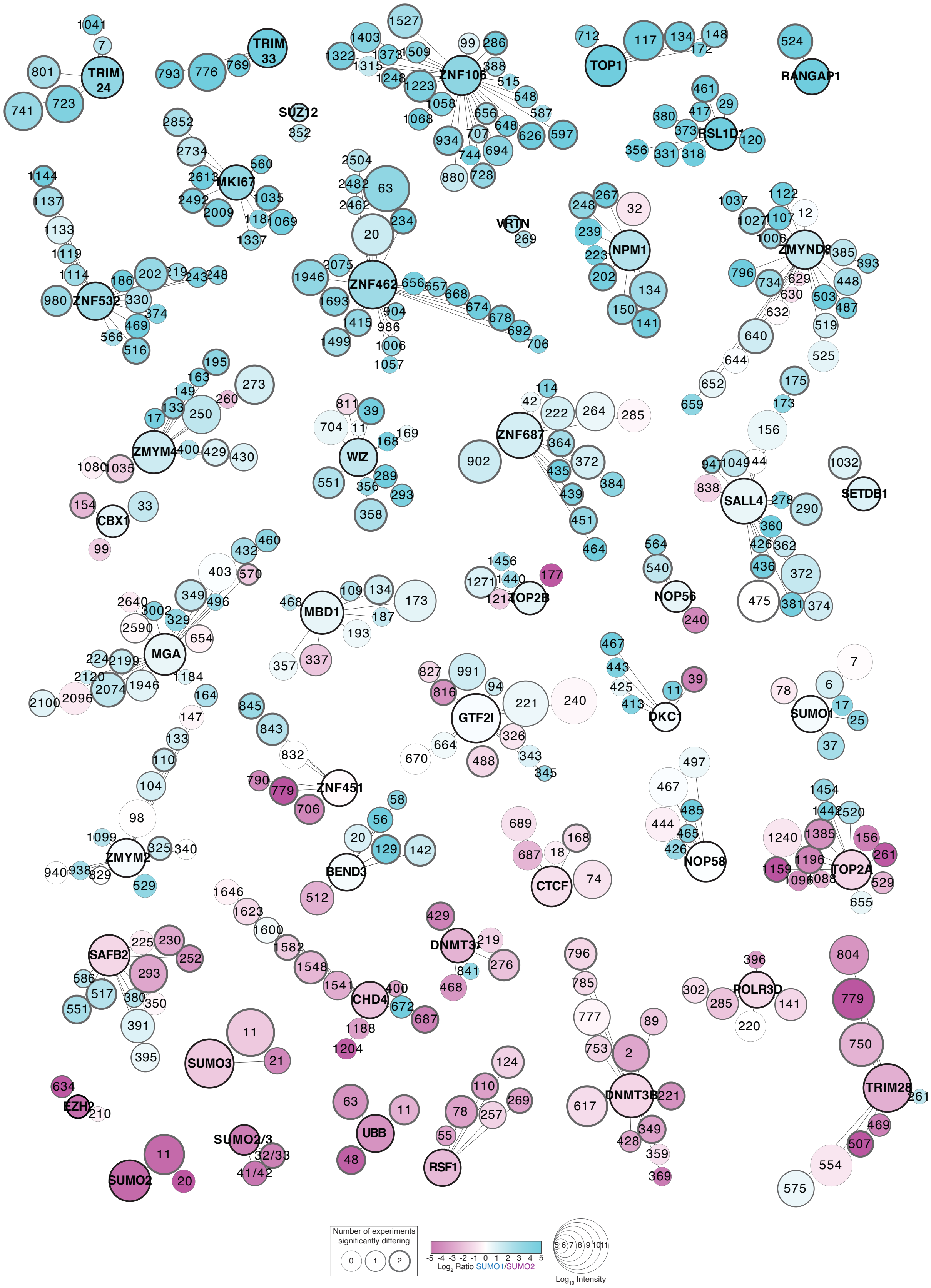

**Supplementary Figure 4. Schematic presentations summarising the SUMO1 and SUMO2 proteomic data for a selection of substrates in hiPSCs.**

Schematic presentations of selected substrates found to be SUMO modified in hiPSCs. Proteins nodes are labelled by name and site nodes by number. SUMO preference is represented by colour considering all GGK peptides (protein nodes) or individual site peptides (site nodes). Peptide intensity is represented by the size of the node (see key). Site node border line thickness represents number of experiments in which it was found to show a significant SUMO preference. Edges linking sites to proteins are positioned relative to their position in the linear protein sequence with first and last residues positioned at the top of the protein node. Substrates are organised from generally SUMO1 preferential (top), to SUMO2 preferential (bottom).

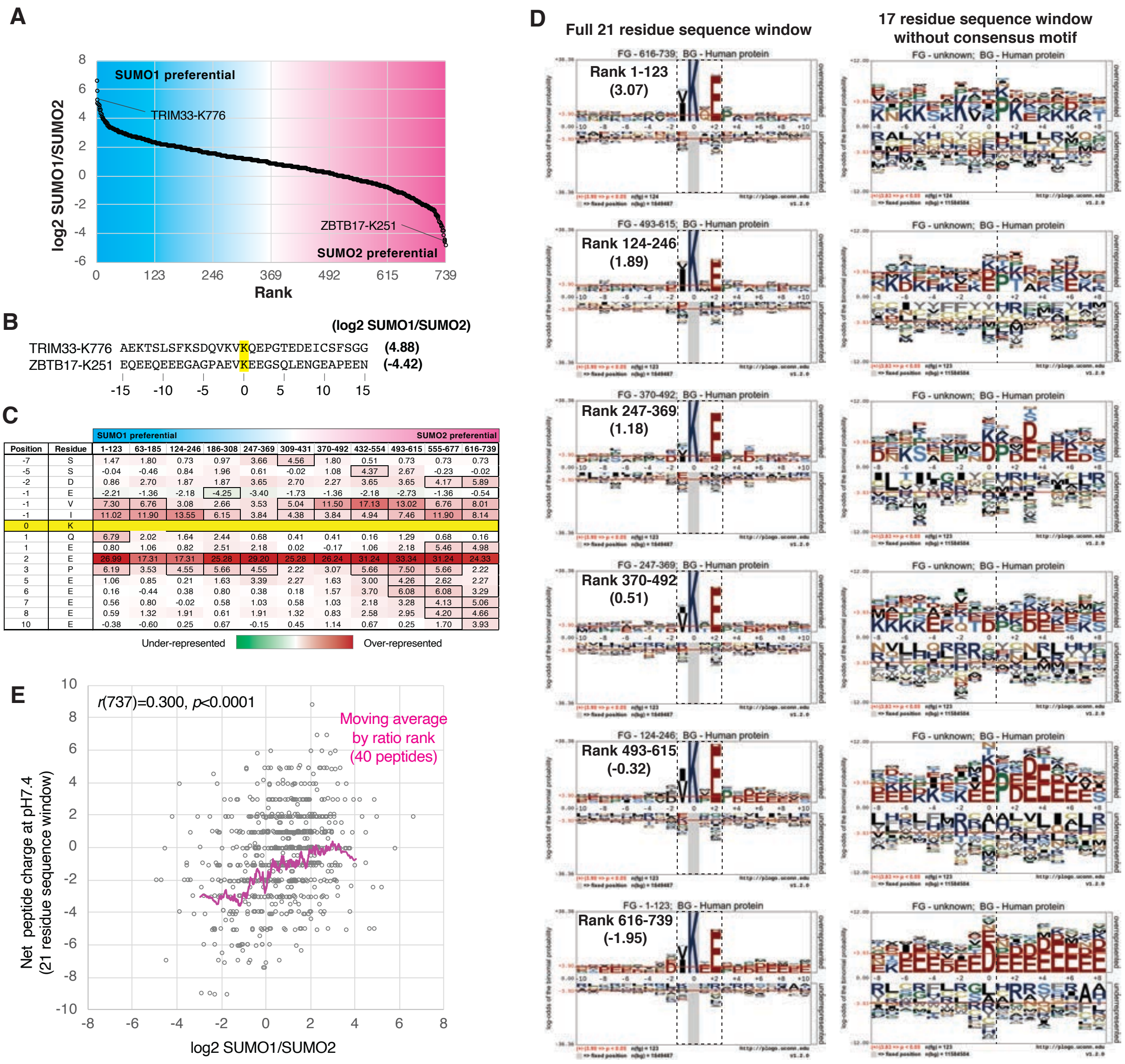


| Position | Residue | 1-123 | 63-185 | 124-246 | 186-308 | 247-369 | 309-431 | 370-492 | 432-554 | 493-615 | 555-677 | 616-739 |
|---|---|---|---|---|---|---|---|---|---|---|---|---|
| -7 | S | 1.47 | 1.80 | 0.73 | 0.97 | 3.66 | 4.56 | 1.80 | 0.51 | 0.73 | 0.73 | 0.73 |
| -5 | S | -0.04 | -0.46 | 0.84 | 1.96 | 0.61 | -0.02 | 1.08 | 4.37 | 2.67 | -0.23 | -0.02 |
| -2 | D | 0.86 | 2.70 | 1.87 | 1.87 | 3.65 | 2.70 | 2.27 | 3.65 | 3.65 | 4.17 | 5.89 |
| -1 | E | -2.21 | -1.36 | -2.18 | -4.25 | -3.40 | -1.73 | -1.36 | -2.18 | -2.73 | -1.36 | -0.54 |
| -1 | V | 7.30 | 6.76 | 3.08 | 2.66 | 3.53 | 5.04 | 11.50 | 17.13 | 13.02 | 6.76 | 8.01 |
| -1 | I | 11.02 | 11.90 | 13.55 | 6.15 | 3.84 | 4.38 | 3.84 | 4.94 | 7.46 | 11.90 | 8.14 |
| 0 | K | | | | | | | | | | | |
| 1 | Q | 6.79 | 2.02 | 1.64 | 2.44 | 0.68 | 0.41 | 0.41 | 0.16 | 1.29 | 0.68 | 0.16 |
| 1 | E | 0.80 | 1.06 | 0.82 | 2.51 | 2.18 | 0.02 | -0.17 | 1.06 | 2.18 | 5.46 | 4.98 |
| 2 | E | 26.99 | 17.31 | 17.31 | 25.28 | 29.20 | 25.28 | 26.24 | 31.24 | 33.34 | 31.24 | 24.33 |
| 3 | P | 6.19 | 3.53 | 4.55 | 5.66 | 4.55 | 2.22 | 3.07 | 5.66 | 7.50 | 5.66 | 2.22 |
| 5 | E | 1.06 | 0.85 | 0.21 | 1.63 | 3.39 | 2.27 | 1.63 | 3.00 | 4.26 | 2.62 | 2.27 |
| 6 | E | 0.16 | -0.44 | 0.38 | 0.80 | 0.38 | 0.18 | 1.57 | 3.70 | 6.08 | 6.08 | 3.29 |
| 7 | E | 0.56 | 0.80 | -0.02 | 0.58 | 1.03 | 0.58 | 1.03 | 2.18 | 3.28 | 4.13 | 5.06 |
| 8 | E | 0.59 | 1.32 | 1.91 | 0.61 | 1.91 | 1.32 | 0.83 | 2.58 | 2.95 | 4.20 | 4.66 |
| 10 | E | -0.38 | -0.60 | 0.25 | 0.67 | -0.15 | 0.45 | 1.14 | 0.67 | 0.25 | 1.70 | 3.93 |

**Supplementary Figure 5. Detailed Sequence logo analysis of SUMO modification sites.**

**A**. Distribution by rank of the log2 SUMO1/SUMO2 ratios for 739 sites. Examples of SUMO1-preferntial (TRIM33 K776) and SUMO2-preferential (ZBTB17 K251) sites are indicated. **B**. 31 residue sequence windows for TRIM33 K776 and ZBTB17 K251. **C**. Residue over- and under-representation within sequence windows of SUMO sites grouped by log2 SUMO1/SUMO2 ratio as shown in A. Only amino-acid positions with at least one significant over or underrepresentation are included. Values are -log10 odds of the binomial probability calculated by pLogo using the human proteome as background. Positive values (red fill) show over-representation and negative values (green fill) show under-representation. Boxes with a black borders are statistically significant ($p$<0.05). **D**. Sequence logos generated by pLogo (ref) for 6 groups of target lysines grouped by log2 SUMO1/SUMO2 as shown in A and used to generate the table in C. Rank range is indicated and average log2 SUMO1/SUMO2 shown in brackets. **E**. Relationship between log2 SUMO1/SUMO2 and net charge at pH 7.4 for the 21 residue sequence window for 739 sites. Pearson correlation is indicated and pink line shows 40 peptide moving average by rank of log2 SUMO1/SUMO2.

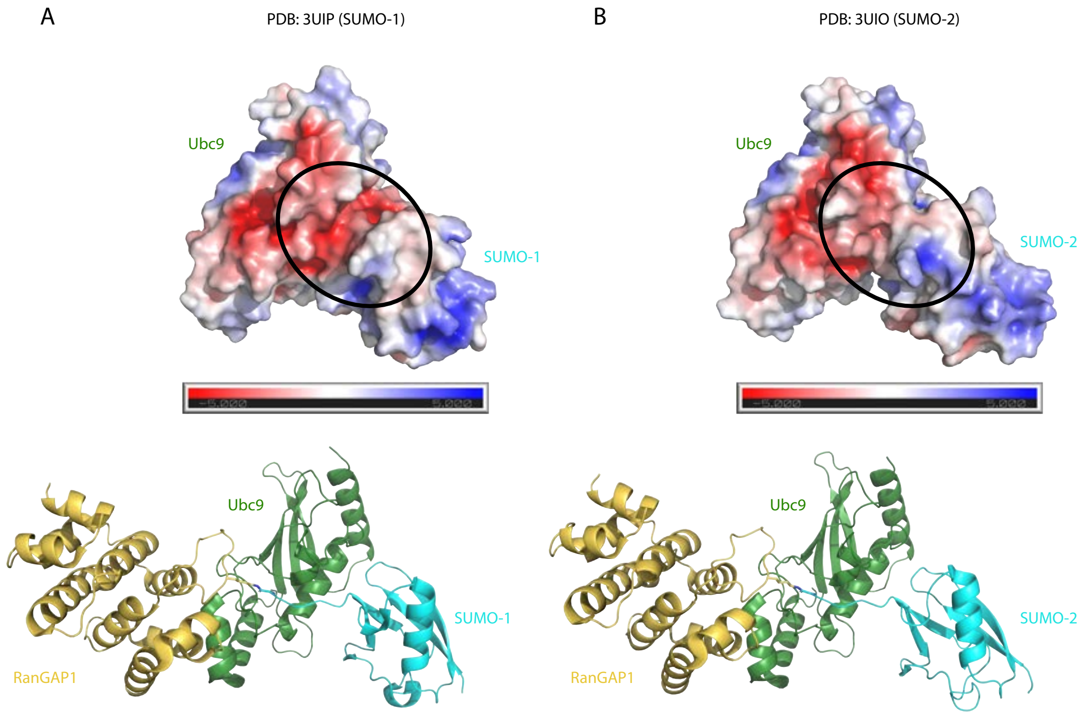


**Supplementary Figure 6. Comparison of the electrostatic potential surface of SUMO-1 and SUMO-2 in complex with Ubc9.**

**A**. Electrostatic potential surface generated at pH 7.0 for Ubc9 and SUMO1 with accompanying electrostatic potential range bar (top panel) and cartoon representation of Ubc9, SUMO1 and RanGAP1 (bottom panel) based on PDB 3UIP. RanGAP1 and RanBP2 have been omitted from the top panel, while RanBP2 has been omitted from the bottom panel, for clarity. A black oval in the top panel highlights the electrostatic potential surface in the active site of Ubc9 and SUMO1, which is presented to substrate. **B.** Same as in A but for PDB 3UIO, which contains SUMO2 instead of SUMO1.
